# Functional mapping of N-terminal residues in the yeast proteome uncovers novel determinants for mitochondrial protein import

**DOI:** 10.1101/2022.08.19.504527

**Authors:** S Nashed, H El Barbry, M Benchouaia, A Dijoux-Maréchal, N Ruiz Gutierrez, L Gaulier, G Chevreux, S Le Crom, B Palancade, F Devaux, E Laine, M Garcia

## Abstract

N-terminal ends of polypeptides are critical for the selective co-translational recruitment of N-terminal modification enzymes. However, it is unknown whether specific N-terminal signatures differentially regulate protein fate according to their cellular functions. In this work, we developed an in-silico approach to detect functional preferences in cellular N-terminomes, and identified in *S. cerevisiae* more than 200 Gene Ontology terms with specific N-terminal signatures. In particular, we discovered that Mitochondrial Targeting Sequences (MTS) show a strong and specific over-representation at position 2 of hydrophobic residues known to define potential substrates of the N-terminal acetyltransferase NatC. We validated mitochondrial precursors as co-translational targets of NatC by selective purification of translating ribosomes, and found that their N-terminal signature is conserved in Saccharomycotina yeasts. Finally, systematic mutagenesis of the position 2 in a prototypal yeast mitochondrial protein confirmed its critical role in mitochondrial protein import. Our work highlights the hydrophobicity of MTS N-terminal residues and their modification by NatC as critical features for the definition of the mitochondrial proteome, providing a molecular explanation for mitochondrial defects observed in yeast or human NatC-depleted cells. Functional mapping of N-terminal residues thus has the potential to support the discovery of novel mechanisms of protein regulation or targeting.

## INTRODUCTION

As soon as they emerge from the ribosomal tunnel, the protein nascent chains recruit factors that will play key roles in their life cycle (1–5). Such recruitment is dependent on the nature of the protein N-terminal amino acid residues. In particular, the specific recognition and co-translational action of several N-terminal modification enzymes is determined by the type of the amino acid residue directly following the initiator methionine, namely the residue at position 2.

For instance, methionine aminopeptidases (MetAPs) will remove the initiator methionine (iMet) in a large fraction of the nascent chains displaying an amino acid with a small radius of gyration (Ala, Cys, Gly, Pro, Ser, Thr, or Val) at position 2 (6). In addition, N-terminal acetyltransferases (NATs) catalyze the irreversible covalent attachment of an acetyl group (CH3CO) to the free α-amino group (NH3^+^) at the protein N-terminus. Three major NATs, namely NatA, NatB and NatC, that acetylate nascent chains in a co-translational manner, are conserved from yeast to human (7). The actual impact of the inactivation of NATs on the N-terminal acetylation of subsets of cellular proteins was measured by various proteomics techniques, eventually providing experimental data on the N-terminal acetylation status for up to 10% of the cellular proteome in yeast and in human (8–11). All these studies led to the conclusion that NatA can acetylate the N-termini beginning with Ala, Cys, Gly, Ser, Thr, or Val after removal of iMet, whereas the other two can acetylate the uncleaved iMet if it is followed by a second specific residue, namely Asn, Asp, Gln, or Glu for NatB, and Ile, Leu, Phe, or Trp for NatC. Based on these substrate specificities together with the observed frequencies of the 20 amino acids at position 2 in proteomes and the partial N-terminal acetylome data, it was estimated that 80−90% of human and 50−70% of yeast proteins could potentially be acetylated by one of these three enzymes (7, 12, 13). Finally, Methyl-, myristoyl-, and palmitoyltransferases can also selectively modify, during or after translation, the N-terminus of proteins after iMet cleavage.

The biological importance of these N-terminal modifications is underlined by the strong defects observed when the corresponding enzymes are inactivated. Complete inactivation of MetAPs activity is lethal in *Escherichia coli*, *Salmonella typhimurium*, and *Saccharomyces cerevisiae*, and the identification of human MetAP1 and MetAP2 as targets for putative anticancer drugs further confirmed the importance of this enzyme family (14, 15). In addition, N-terminal acetylation has been implicated in several diseases, including cancers, developmental disorders, as well as Parkinson's disease (5). At the molecular level, the importance of the nature of the second residue and the associated N-terminal modifications have been implicated in all major steps of proteins life cycle including folding and aggregation, protein interactions and complex formation, protein subcellular targeting, and protein turnover through proteasomal degradation pathways (reviewed in [6]). In particular, during the last decades, the exploration of the N-end rule pathway, linking the nature of the N-terminus to protein *in vivo* stability has emphasized the importance of protein N-end signatures and demonstrated that iMet cleavage and N-terminal acetylation are important players in the early control of proteins life cycle (16–19). Finally, preferences for the use of amino acids at position 2, different from those observed globally in the proteome, have been described in various species, suggesting that selection pressures act on this particular protein position. In addition, these biases are taxon-specific, indicating progressive changes during evolution (20).

This extensive knowledge on the general importance of protein N-terminal signatures sharply contrasts with the lack of data describing their potential implication in specific functional pathways. To address this issue, we have developed an in-silico approach at the proteome scale allowing for the systematic detection of functional biases in the use of the 20 amino acids at position 2 of proteins. The rationale of our approach is that, given the importance of the residue at position 2 in the life cycle of proteins, evolutionary constraints could have led to the selection of specific amino acids at this position in groups of proteins from the same cellular compartment or involved in the same biological pathway. As a proof of concept, we analyzed the S. *cerevisiae* proteome to statistically assess all significant and specific overrepresentations of one or more amino acids at position 2 in protein subsets sharing the same Gene Ontology (GO) annotations. Using this approach, we were able to identify various groups of proteins with common GO annotations and characterized by particular amino acid preferences at position 2. We hypothesized that these newly identified N-terminal signatures are likely associated with important functional roles. We further characterized, with a combination of *in-silico* analyses and experimental assays, the function of the strong position-specific bias detected at position 2 for mitochondrial proteins.

Most mitochondrial proteins are translated by cytoplasmic ribosomes and addressed to the mitochondrial compartment via a cleavable presequence, the so-called Mitochondrial Targeting Sequence (MTS). We found that, throughout the Saccharomycotina yeast lineage, MTS-bearing mitochondrial precursors present, just after the iMet, predominantly hydrophobic amino acids known to define potential NatC substrates (i.e., Leu, Phe, Ile, and Trp). Affinity-selective purification of the translating ribosomes then confirmed their co-translational recognition by this N-terminal acetyltransferase. Finally, we demonstrated the functional significance of the bias detected at position 2 of MTSs by site-directed mutagenesis of this position in a dominant negative allele of the essential mitochondrial protein Hsp60p. Only the amino-acids found to be overrepresented at position 2 of MTSs in our *in-silico* analyses allowed the efficient mitochondrial import of the toxic Hsp60p, as revealed by the associated loss of cell viability.

Our work has revealed an unknown and critical feature of MTSs. Defects in mitochondrial biogenesis are implicated in a number of serious pathologies, including neuropathies, cardiovascular disorders, myopathies, neurodegenerative diseases and cancers (21). In a long-term perspective, understanding the role of this signature for mitochondrial protein import is therefore crucial for human health. For example, this knowledge could be very useful for the design of mitochondrial gene therapies based on the targeting of specific proteins to the mitochondrial compartment (22). Beyond the results obtained for the proteins of the mitochondrial compartment, we provide the community with the first functional mapping of N-terminal residues, whose future exploration will undoubtedly contribute to the discovery of novel mechanisms of protein regulation or targeting.

## RESULTS

### Detection of functional biases in amino acid usage at position 2 in *S. cerevisiae* proteome

The amino acid usage at position 2 of *S. cerevisiae* proteins (Fig. 1a) is clearly different from the average amino acid distribution in its proteome. Strikingly, we observed a serine at this position in nearly 25% of the yeast proteins. This overrepresentation is both highly significant (HGT score equal to 268 corresponding to a p-value after hypergeometric test equal to 10^−268^, see Methods for a definition of HGT score) and specific to position 2 (aspecificity score of 0%, see Methods). By contrast, several amino acids are underrepresented (HGT score <−3 and aspecificity score <3%), including the hydrophobic residues leucine, isoleucine, and tyrosine as well as charged or polar residues such as arginine, glutamate and glutamine. Such asymmetric distribution, with the over-representation of serine, is conserved in budding yeasts (Fig. Sup1), as indicated by proteome analysis of 17 yeasts spanning nearly 400 million years of evolution of the Saccharomycotina lineage (23, 24).

**Figure 1:**
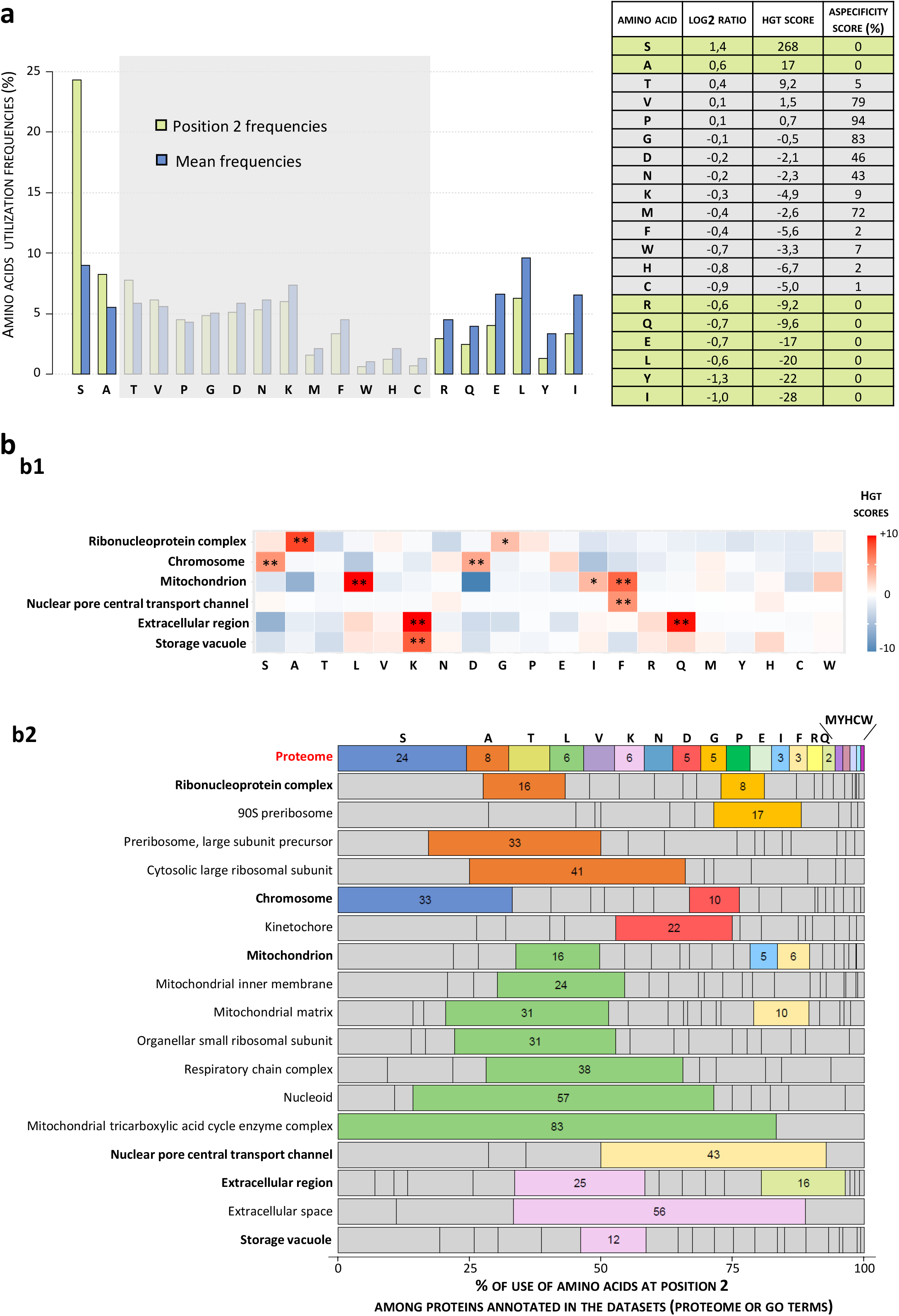
Amino acid usage biases at position 2 of *S. cerevisiae* proteins. (a)Amino acid usage at position 2 compared to their average usage at any position in *S. cerevisiae* proteome. Frequencies (Barplot, left) of the 20 amino acids measured at position 2 among the 5,959 sequences of *S. cerevisiae* proteins retrieved from UniProtKB database (68). Table (right) recapitulates quantitative values calculated to identify significant amino acid usage biases at position 2: position2/mean log2Ratio, HGT score obtained from hypergeometric tests, and aspecificity scores indicating the percentage of positions in sequences with usage bias as high as that measured at position 2 (see Methods). Amino acids for which the absolute value of the HGT score is greater than 3 (p-value<10^−3^) and the aspecificity score is lower than 5% are highlighted in yellow. Such values indicate a significant and specific bias of usage at position 2 revealing a possible selection pressure. (b) Detection of biases in the use of amino acids at position 2 in subsets of proteins of the various cellular compartments of *S. cerevisiae*. We performed a systematic detection of the significant over-representation of amino acids at position 2 among groups of proteins sharing the same GO annotations (see Methods). (b1) Heatmap of HGT scores for the 20 amino acids in GO terms in the “component” category identified with this strategy. Seventy-two GO terms were first identified with HGT score higher than 4 (i.e. p-value<10^−4^) for at least one amino acid. Our reduction algorithm finally retained 6 bestN GO terms corresponding to the minimal number of terms maximizing the number of proteins with the detected usage biases (see Methods). Stars indicate significant HGT scores (*HGT>3, **HGT>4). (b2) Barplot showing the percentages of use of the 20 amino acids at position 2 in retained GO terms compared to their use at the same position in the proteome. For a better characterization of position 2 functional biases, bestF GO terms were retrieved by our algorithm (see Methods) and added to bestN GO terms in bold. Each of them contains proteins included in one BestN GO term and has higher position 2 frequency biases. For each GO term, highlighted amino acids are those for which the percentage of use at position 2 is significantly higher than that observed at the same position in the proteome (HGT score > 3). In figures b1 and b2, the amino acids are sorted in decreasing order of use at position 2 in the proteome.

We further developed a computational approach to explore the functional relevance of amino acid biases at the proteome scale and assess their statistical significance. We took advantage of the robust Gene Ontology annotations of the *S. cerevisiae* proteome to detect all significant and specific overrepresentations of one or more amino acids at position 2 in protein subsets defined by common GO annotations.

In a first step, we scanned the diverse GO terms describing the components, the cellular pathways or the protein molecular functions and we calculated the frequencies of the 20 amino acids at position 2 in the associated protein subsets. We then identified the subsets displaying significantly higher frequencies than expected by the overall distribution of amino acids at position 2. We assessed the significance of the enrichments by calculating the HGT scores derived from p-values obtained in hypergeometric tests. With this strategy, we identified 232 GO terms with a significantly increased frequency (HGT>4) for at least one amino acid. These GO terms were distributed as follows in the different GO categories: 72 GO terms corresponding to components, 138 to cellular pathways and 22 to molecular functions (Table S1). We designed a dedicated algorithm to reduce this initial list and eliminate information redundancy in each category. Rather than relying on the hierarchical relationships between the GO terms, as usually done by existing solutions (25), our algorithm directly analyzes the characteristics of the protein subsets defined by these GO terms, such as the number of proteins, the enrichment factors, and the overlaps between the subsets. In a first filtering step, we eliminate GO terms with the lowest frequency bias (<1.8 fold change) and corresponding to very generic components or processes, which typically include a very high number of proteins. Our algorithm then retains the GO terms encompassing the largest number of proteins and maximizing the coverage of the original dataset (called BestN GO terms, see Methods). When possible, it complements this set of bestN GO terms with the one or several smaller GO terms displaying the highest position 2 frequency biases (called BestF GO terms, see Methods). Each BestF is ultimately linked to the nearest BestN based on the overlap of their respective set of proteins.

Figure 1b illustrates the results of this filtering for the GO category “components” and represents the amino acid usage biases observed in the 17 GO terms that were finally retained after processing the initial list of 72 GO terms. Results for the other two GO categories, “cellular pathways” and “molecular functions”, are available as supplementary data (see Table S1 and Fig. sup2 in which the original lists were reduced from 138 to 21 and from 22 to 10 GO terms, respectively). The final list of GO terms “components” includes the 6 BestN GO terms “Ribonucleoprotein complex”, “Chromosome”, “Mitochondrion”, “Nuclear pore central transport channel”, “Extracellular Region” and “Storage vacuole” (fig. 1b1). These GO terms covered 85% of the proteins with the amino acid preferences associated with the 55 GO terms retained after the first filtering step that removed low bias generic GO terms. The 11 GO BestF terms (Fig. 1b2) provided a more detailed picture of protein subsets with specific N-terminal amino acid usage. For example, “90S preribosome” (3.4-fold increase in glycine utilization at position 2) and “large cytosolic ribosomal subunit” (5.1-fold increase in alanine utilization at position 2) led to a much better characterization of the proteins involved in the bias observed in the more general GO term “ribonucleoprotein complex”. Similarly, the “Kinetochore” proteins (4.4-fold increase in aspartate utilization at position 2) explain a large part of the aspartate bias detected in the GO term “Chromosome” since it accounts for 12 of the 21 proteins responsible for this bias detection. The BestF GO terms also pointed to several mitochondrial sub compartments with high preferences for leucine at position 2 (4.0- to 13.8-fold increase in leucine usage), suggesting that this bias involved only a specific subset of the mitochondrial proteins localized in the "Mitochondrial inner membrane” and the “Mitochondrial matrix”.

The physiological significance of the amino acid utilization biases detected remains to be elucidated, and some of them might be related to the activity of N-terminal modifying enzymes such as MetAPs or NATs. One of the strongest biases revealed by our approach is the dramatic overrepresentation of leucine at position 2 in GO terms related to the mitochondrial compartment. In this work, we sought to clarify, in *S. cerevisiae*, the functional significance of usage bias at position 2 of mitochondrial precursors and its potential relationship with NatC.

### The N-terminal Mitochondrial Targeting Sequence has a specific conserved signature at position 2 typical of NatC potential substrates

We first investigated whether the high overrepresentation of leucine at position 2 of the mitochondrial precursors could correlate with other specific features in the amino acid composition of their N-terminal region. Especially, precursors of matrix, inner membrane and intermembrane space proteins are dependent for their mitochondrial import on an N-terminal sequence forming an amphiphilic alpha helix, 25−35 residues-long, enriched in hydrophobic and positively charged residues (26, 27). Such a mitochondrial targeting sequence, called MTS, is observed in 361 of the 726 yeast proteins annotated as mitochondrial with high confidence (Table S2, see Methods). Consistently, the 361 mitochondrial precursors harboring a MTS showed a dramatic overrepresentation of arginine and an underrepresentation of negatively charged residues in the positions 3 to 20 of their N-terminal region. This profile contrasted strongly with that of mitochondrial precursors lacking MTS (Fig2a). Strikingly, we found that the bias at position 2 described above for the proteins associated with mitochondrial GO terms was specific of mitochondrial proteins with a MTS (Fig2a). More precisely, four hydrophobic residues (Leu, Phe, Trp and Ile) are significantly overrepresented (HGT value ranging from 3.9 for isoleucine to 71 for leucine) at position 2 of proteins with MTS (Fig.2a). Furthermore, it should be noted that these biases in favor of Leu, Phe, Trp and Ile are strictly restricted to position 2 of the MTS.

**Figure 2:**
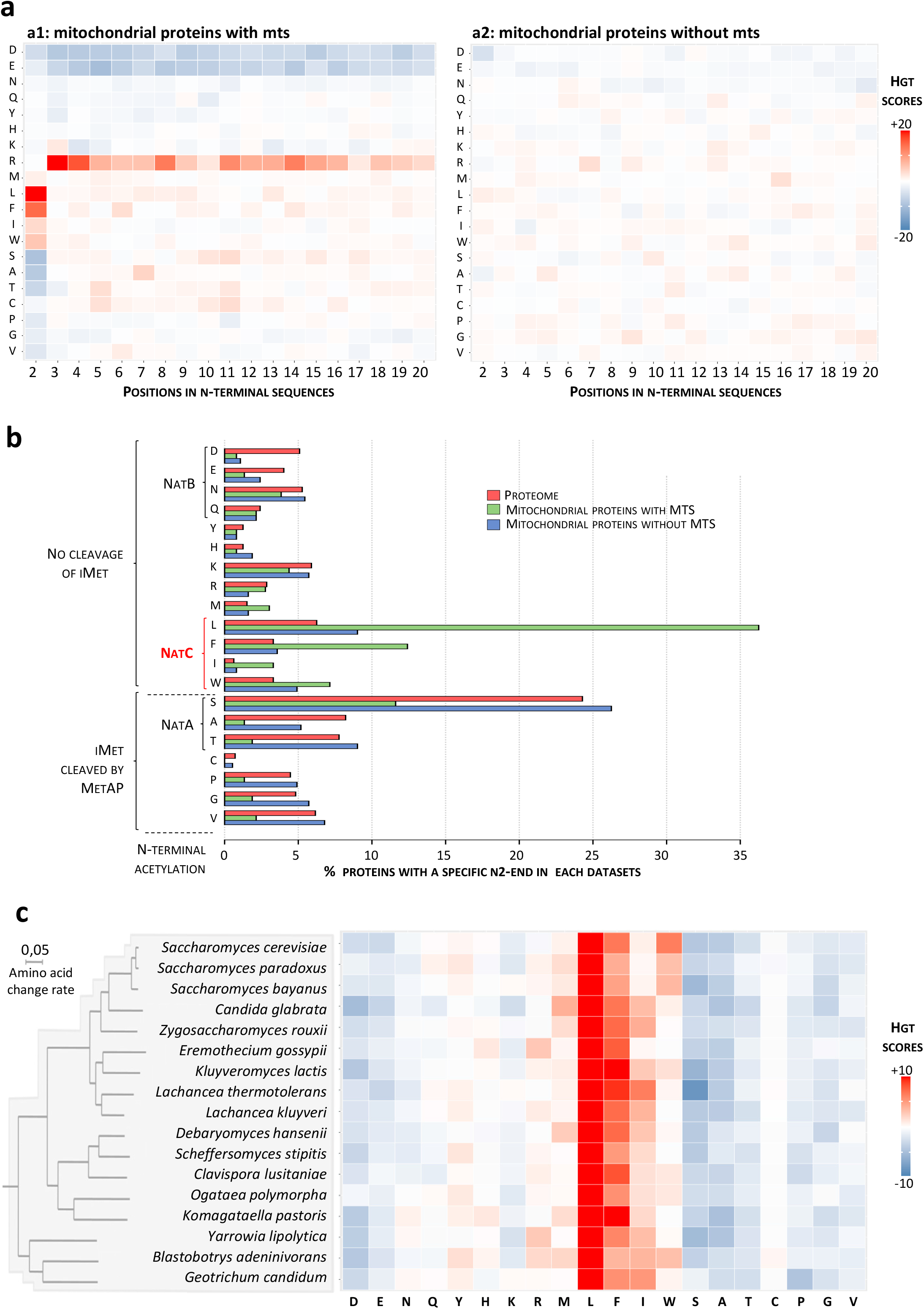
Mitochondrial precursors harboring a MTS are putative substrates of NatC. To characterize the bias detected at position 2 of mitochondrial precursors, the list of mitochondrial proteins extracted from the Gene Ontology database was manually curated (see Methods), and mitochondrial precursors were sorted according to whether they carry a MTS based on annotations retrieved from the Uniprot database (68). The two classes of mitochondrial proteins were then analyzed separately. (a) Heatmap showing the hypergeometric enrichment scores (HGT score) obtained by comparing the amino acid usage at the first 20 positions of the mitochondrial precursors harboring (a1) or not (a2) a MTS with their average usage in the proteome at these same positions. Mitochondrial precursors with a MTS display the expected MTS characteristics (positive bias toward R and negative bias toward D and E) and exhibit specific biases at position 2 (positive bias toward F, L, I and W). This position 2 specific biases were not observed among precursors without N-terminal addressing sequence. (b) Distribution of the 20 amino acids at position 2 in the proteome and among the two classes of mitochondrial precursors. Several amino acids are overrepresented at position 2 of mitochondrial precursors harboring a MTS, which is not the case for the mitochondrial precursors lacking a MTS. Amino acids showing significant increase compared to the proteome are those that define NatC putative substrates according to the literature: Leucine (6 fold increase: 36% vs 6%, p-value = 10^−71^), Phenylalanine (4 fold increase: 13% vs 3%, p-value = 10^−15^), Tryptophan (5 fold increase : 3% vs 0.6%, p-value = 10^−6^) and Isoleucine (2 fold increase: 7% vs 3%, p-value = 10^−4^). (c) Heatmap showing, for 17 budding yeasts of the Saccharomycotina lineage, the HGT score obtained by comparing amino acids usage at position 2 of MTS-harboring mitochondrial precursors and usage at the same position in the proteome. The positive bias, particularly for leucine, at position 2 of mitochondrial precursors is conserved in all analyzed species. The phylogenic tree was retrieved and modified from (23).

Quantitative analysis of the distribution of amino acids at position 2 of the MTSs (Fig. 2b) shows the importance of this previously unknown N-terminal signature of the mitochondrial addressing sequence: nearly 60% of them have an N-end signature with a Leu, Phe, Trp and Ile, which contrasts strongly with the low representation of the latter at position 2 in the proteome (see L, F, I, W residues frequencies in Fig. 1a). For instance, leucine accounts for more than 35% of the amino acids observed at position 2 of the MTSs, whereas it is poorly used in the *S. cerevisiae* proteome at this position (Fig 1a). Interestingly, we also found an overall significant under-representation at position 2 of the MTSs of amino acids with small radii of gyration (Gly, Ala, Ser, Cys, Thr, Pro and Val), known to induce the cleavage of iMet by methionine aminopeptidase (6). Finally, the analysis of amino acid biases in the MTSs of 17 budding yeasts of the Saccharomycotina lineage (23, 24), demonstrated that this specific signature at position 2 is a conserved characteristic of mitochondrial addressing sequences (Fig.2c). Indeed, in all the studied species, we observed: (1) a strong over-representation of the same hydrophobic amino acids at position 2 and in particular of leucine whose HGT score exceeds the value of 20 in all species, and (2) a bias restricted to position 2 since the HGT scores of leucine, phenylalanine, isoleucine and tryptophan were systematically lower than 3.9 for all the other positions of the MTSs (Fig. Sup1). Interestingly, the conserved amino-acid bias described above at position 2 of MTSs perfectly correlated with the known specificities of N-acetyl transferases in yeast (Fig. 2b). Indeed, leucine, phenylalanine, isoleucine and tryptophan, which are overrepresented at this position, are known to define potential substrates of NatC in yeast when positioned after iMet (9). Conversely, serine, alanine and threonine, which are the most under-represented at this position, are known to induce iMet cleavage and subsequent targeting by NatA (8).

Together, these observations strongly suggest that position 2 of N-terminal mitochondrial targeting sequences is under selective pressure and that the nature of the residue at position 2 is crucial for the fate of mitochondrial precursors imported through the MTS-dependent pathway. The correspondence between the amino acid bias at position 2 of MTSs and the preferences of NatC suggests a link between mitochondrial precursors fate and NatC targeting. They raised two important questions addressed in the following sections. First, are mitochondrial precursors indeed targeted by the N-terminal acetyl transferase NatC? And second, what are the functional consequences of changing the residue located at position 2 of a typical MTS?

### Selective Translating Ribosome Affinity Purification reveals co-translational targeting by NatC of mitochondrial precursors harboring the identified N-terminal signature

Our current knowledge of the substrate specificities of NatC in yeast identifies proteins with ML-, MF-, MI- and MW-N-termini as its potential substrates. It was deduced from the observation of the loss of N-terminal acetylation of various iso-1 cytochrome c reporter proteins mutated at position 2 in yeast strains lacking NatC, which was further confirmed for a small subset of cellular proteins (9, 28–30). Partial N-terminal acetylome of *S. cerevisiae* were more recently obtained (8, 10, 11) that covered up to 10% of the proteome, corresponding to the most abundant cellular proteins. Unfortunately, these data included very few NatC potential substrates (between 0 and 20 for each NatC typical N-termini) and, to date, there are no published data describing globally the *in vivo* substrates of NatC by studying the impact of NatC depletion on the yeast N-terminal acetylome.

Mitochondrial precursors steady state levels are extremely low, since, after their synthesis, they are quickly imported in mitochondria where their N-terminal addressing sequence is cleaved (26). Hence, detecting them with protein-based approaches is challenging. To overcome this limitation, we decided to use the Selective Affinity Purification of Translating Ribosomes (sel-TRAP) method, to capture by immunoprecipitation the various NatC-targeted nascent chains, as well as their associated mRNAs, which we characterized using transcriptomic analyses (Fig. 3a, see also microarray data in Table S3). This approach had already proven sufficient sensitivity to study the selectivity of various nascent chain associated factors (31–34). To validate our strategy, we also conducted sel-TRAP experiments on NatA and NatB, which preferences were previously confirmed by the impact of their deletion on N-terminal acetylome. Hence, we immunoprecipitated protein A-tagged versions of the catalytic subunits of each of the three NATs, Ard1p (NatA), Nat3p (NatB), and Mak3p (NatC) (Fig. 3b, top and middle panel). The detection of rRNAs in the immunoprecipitated fractions (Fig. 3b, bottom panel, IP lane) and the analysis of the protein fractions of the three immunoprecipitations compared with the mock experiment confirmed that we had purified translating ribosomes that were mostly distinct by the specific presence of the catalytic and accessory subunits of the immunoprecipitated N-terminal acetyltransferase complexes (Fig. 3c, see also proteomic data in Table S4). Transcriptomic analyses of the immunoprecipitated samples compared to the input RNAs identified non-overlapping sets of enriched mRNAs likely encoding the respective substrates of each NAT. As a control, we also analyzed mRNA co-immunoprecipitated with the canonical Rpl16a ribosomal protein, which confirmed the specificity of the mRNAs identified in the NATs sel-TRAP experiments (Fig. 3c).

**Figure 3:**
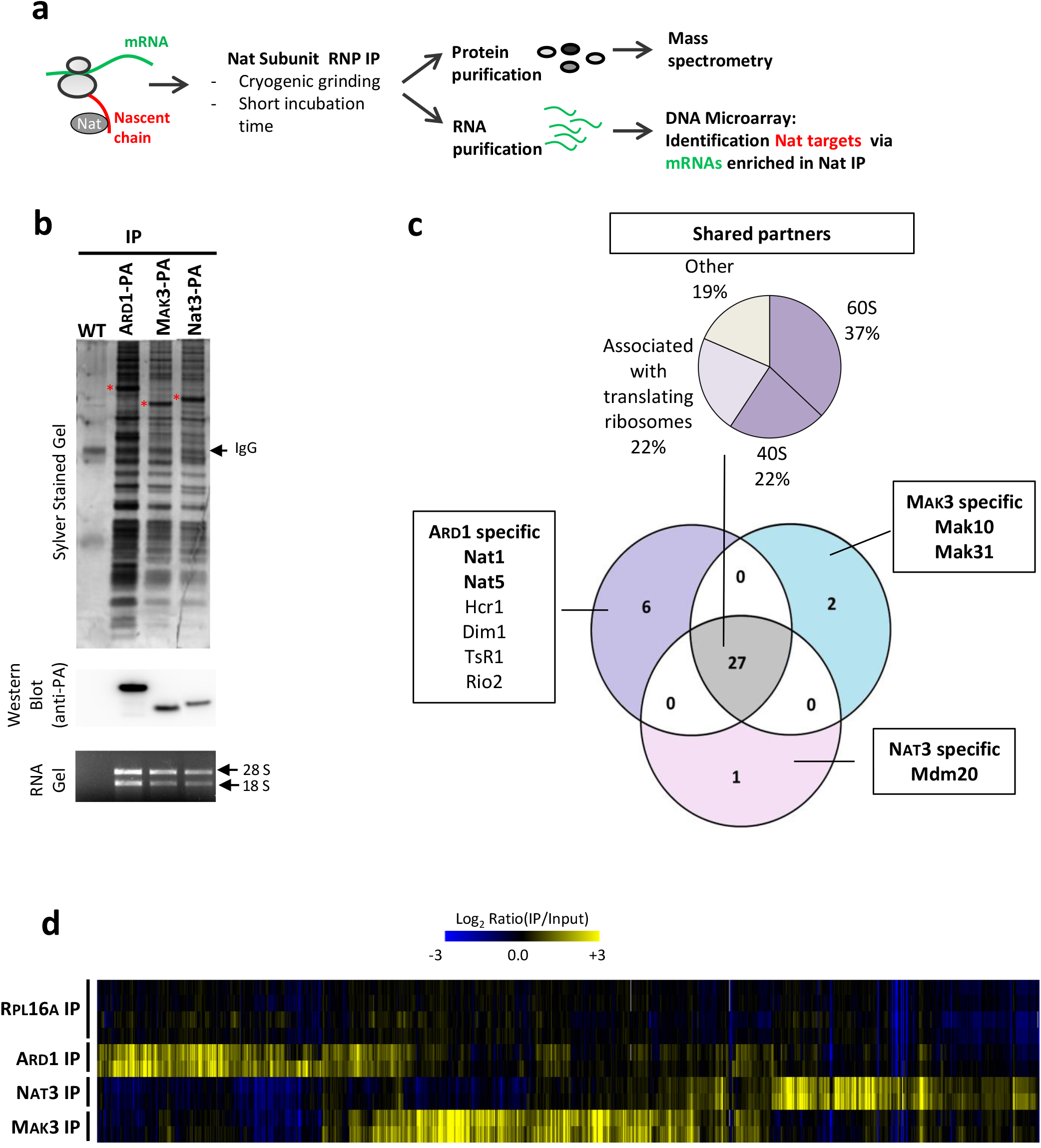
Selective Translating Ribosome Affinity Purification (sel-TRAP): a comprehensive approach to identify translational targets of N-acetyltransferases. (a) Experimental approach. Immunoprecipitations of NatA, NatB and NatC were performed on cells expressing respectively Ard1-PA, Nat3-PA and Mak3-PA. A control experiment was also performed with an untagged strain (WT). Protein (b and c) and RNA (b and d) contents were analyzed from immunoprecipitated samples (IP). (b) Top panel: Immunoprecipitated fractions obtained from untagged or PA fusion protein expressing-cells were analyzed by SDS-PAGE followed by silver staining. The position of the Ard1-PA, Nat3-PA and Mak3-PA fusion and IgG are indicated. Middle panel: The same samples were analyzed by SDS-PAGE followed by western-blots using antibodies recognizing the PA epitope. Bottom panel: Following RNA isolation, immunoprecipitated fractions obtained were analyzed by agarose gel electrophoresis with ethidium bromide staining. The bands corresponding to 28S and 18S rRNA are indicated. (c) Comparison of the composition of NatA (Ard1), NatB (Nat3) and NatC (Mak3)-associated ribosomes. Immunoprecipitations of proteinA-tagged versions of the catalytic subunit of the three Nats were followed by mass spectrometry analysis of their protein partners. Venn Diagram represents the overlap of the three different sets of co-immunoprecipitated proteins: for each sector the number and the name of interactants reproducibly detected in two independent experiments are indicated (Table S4). Partners indicated in bold correspond to the known subunits of the N-terminal acetyltransferase complexes. Chart pie shows the functional distribution of proteins for shared partners in the three datasets. (d) To identify mRNAs specifically enriched after sel-TRAP with cells expressing protein A tagged versions of Ard1, Nat3 or Mak3, RNA contents from total extract (Input) and immunoprecipitated samples (IP) were analyzed using microarray technology. As a control, the same approach was applied with the whole set of ribosomes immunoprecipitated using cells expressing a proteinA-tagged version of Rpl16a. Hierarchically clustered heat map represents IP/input log2(ratios) for mRNAs enriched, in two independent experiments, in at least one of the NAT immunoprecipitations compared to Rpl16a (Table S3). Each line represents an experiment, each column a gene.

To verify that the NATs sel-TRAPs have specifically purified their respective substrates, we performed a threshold-independent analysis of the proportion of amino acids encoded at ORFs position 2 in the transcriptomic data. We detected a specific and significant increase in serine in the NatA immunoprecipitation experiment, aspartate in that of NatB, and leucine and phenylalanine in that of NatC (Fig. Sup3). These results are globally consistent with the known substrate specificity of the three NATs in *S. cerevisiae* (see Introduction). However, all the expected N-termini were not detected as enriched, which could reflect differences in the efficiency of recognition of their potential substrates by the NATs. Consistent with this idea, N-terminal acetylome data has previously revealed that, in many cases, only a fraction of NATs expected substrates is actually acetylated (8, 10, 11). For example, among the proteins potentially targeted by NatA, almost all of those with a Ser- N-termini were acetylated, whereas only about 50% of those with Ala- or Thr-N-termini were, and almost none of the other potential NatA targets (Pro-, Gly-, and Val- N-termini). Similarly, most of the few NatC potential substrates detected in N-terminal acetylome data were only partially acetylated. It should also be kept in mind that sel-TRAP technology captures co-translational targets that co-purify with the immunoprecipitated NATs and that the transient nature of enzyme-substrate connections could limit the detection of the weakest interactions.

As N-terminal acetylome data in absence of NatC are not available, sel-TRAP allowed for the first time a global study of NatC targets *in vivo* in *S. cerevisiae*. After immunoprecipitation of Mak3p, we observed in our threshold-independent analysis a significant increase specific to leucine and phenylalanine at position 2 of the proteins encoded by the most enriched mRNAs (green and red curves in Fig.4a1, see also Fig. Sup3). The significance of these increases was confirmed by the observation that the HGT scores for the ML- and MF-N-termini reached a maximum value near 15 and 25, respectively (green and red curves in Fig.4a2). This is consistent with the known substrate specificity of NatC and suggests more efficient co-translational acetylation of the initiator methionine when followed by these residues rather than by isoleucine or tryptophan. Furthermore, in agreement with our prediction, nearly 50% of the enriched mRNAs (enrichment threshold value equal to 0.8) encoded mitochondrial precursors (Fig 4b) and only those imported through the MTS-dependent pathway were observed enriched in the sel-TRAP data (Fig 4c). Among these precursors, the most significant enrichments were observed for those with a leucine (HGT=24), phenylalanine (HGT=23), or isoleucine (HGT=5) at position 2 of their mitochondrial addressing sequence (Fig 4d).

**Figure 4:**
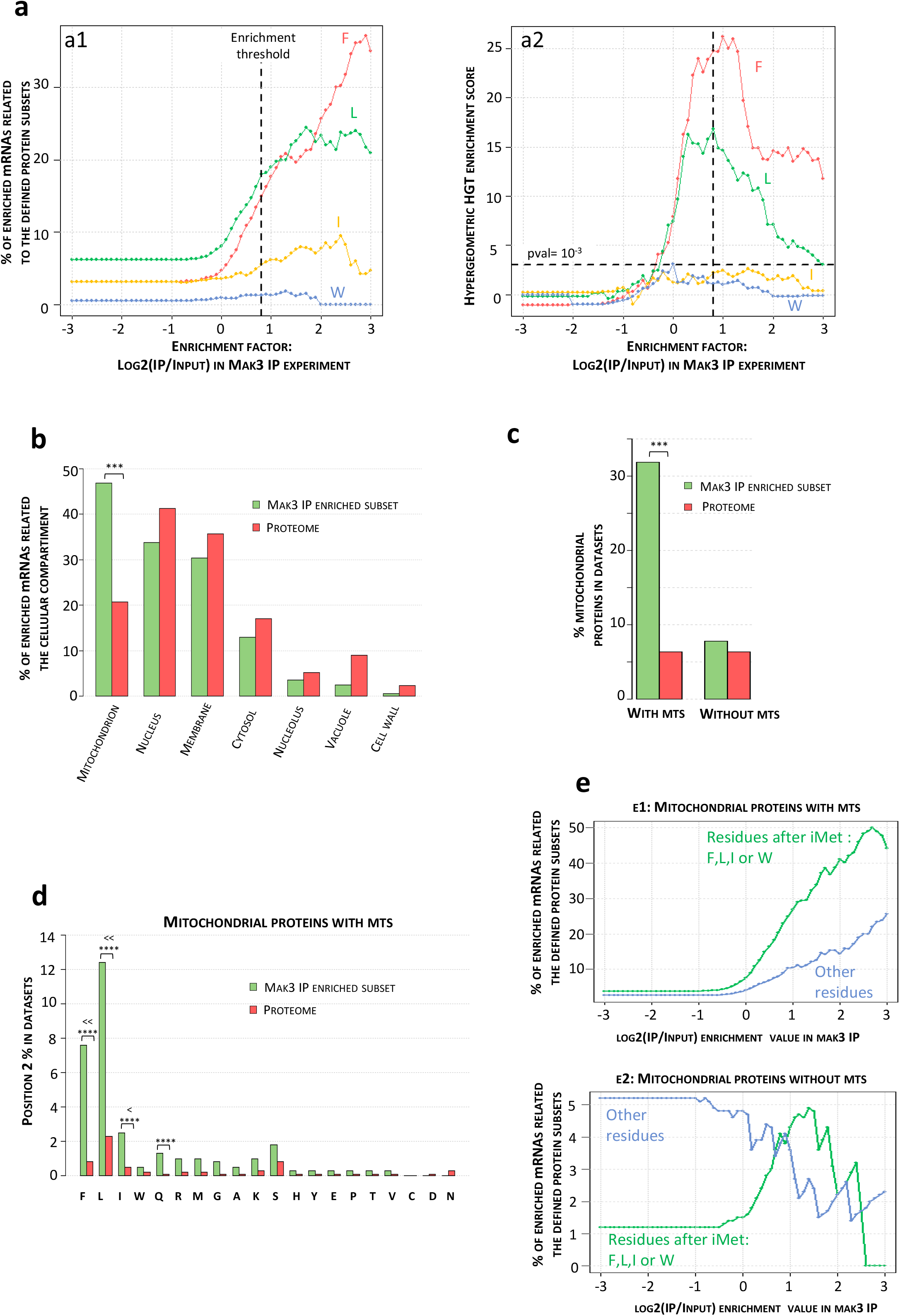
Analysis of NatC selectivity with sel-TRAP technology confirms mitochondrial precursors harboring a MTS as its main targets. Global analysis of mRNAs copurified after immunoprecipitations (IP) from cells expressing Mak3-PA allowed the indirect characterization of nascent protein chains translated by ribosomes associated with NatC. For each cellular protein, the corresponding mRNA enrichment value, defined as IP/input log2(ratio), was calculated after microarray hybridizations of cDNA reverse-transcribed from immunoprecipitation (IP) and input samples. Since NatC, as well as other NATs, is known to target proteins according to the nature of their N-terminus, the evolution of the proportion of the 20 possible residues at position 2 associated with the subsets of mRNAs detected with increasing enrichment values in IP experiments was studied (Fig. sup3). (a) Curves obtained for N-terminal residues defining the putative substrates of NatC (i.e. F, L, I and W), are shown in a1. To assess the statistical significance of the observed increases, the evolution of HGT scores as a function of the enrichment value is also plotted (a2), which allowed the detection of a significant enrichment of the MF- and ML- N-termini in the most enriched protein set (horizontal dotted line indicates the HGT score cutoff of 3 corresponding to a p-value equal to 10^−3^). Because highest HGT scores are reached for an enrichment value around 0.8, this value was chosen as a threshold (vertical dotted lines in a1 and a2) to determine enriched mRNAs identified with sel-TRAP experiments of NatC. (b) The percentages of each residue at position 2 among the identified targets (enrichment value >= 0.8) were compared to those observed in the proteome. MF- and ML- N-termini represent respectively 15 and 18% of the enriched dataset versus 3 and 6% in the proteome. (c) The percentages, among the identified targets, of proteins with GO annotation corresponding to the main cellular compartments were compared to their respective percentage in the proteome. This analysis confirmed the strong selectivity of NatC for mitochondrial precursors, which represent 47% of the identified NatC targets compared to 21% in overall cellular proteins (p-value = 10^−33^). (d) The percentages of the two classes of mitochondrial precursors among the identified targets (enrichment value >= 0.8) were compared to those observed in the proteome. Mitochondrial precursors with a MTS represent 32% of the identified NatC targets versus 6% in the proteome (p-value=10^−54^). No significant enrichment of the mitochondrial precursors without MTS was detected in the sel-TRAP data. (e) The percentages of the different N-termini among the MTS identified as NatC targets were compared with those observed globally among MTSs. Significant enrichment was detected for the MF-, ML-, and MI- N-termini (p-values equal to 10^−24^, 10^−23^, and 10^−25^, respectively) and to a lesser extent for the MQ- N-termini (p-value equal to 10^−4^). Stars indicate significant p values (p value=<10^−4^). (f) Evolution of the proportion of specific N-termini in the two classes of mitochondrial precursors in the mRNA subsets defined with increasing enrichment values. Precursors with (e1) or without mts (e2) were sorted depending on whether they carry at position 2 a putative NatC-targeted residue (L, F, W or I). As expected, we observed a significant increase in the proportion of precursors with NatC-targeted N-termini, regardless of whether they carry a MTS. This analysis also revealed an increase in the proportion of precursors with MTSs that lack an N-terminus recognized by NatC, suggesting that NatC may recognize other MTS features.

Intriguingly, we also observed in the sel-TRAP data a slight enrichment of MTSs without a typical signature of NatC targets at position 2 (amino acid other than F, L, I, W in Fig. 4d). With the exception of glutamine at position 2 (HGT=4), these enrichments were not significant, but this could be due to the low number of precursors with a given amino acid at position 2 of their MTSs sequence (between 3 and 42). Indeed, a threshold-independent analysis showed that the enrichment of mitochondrial precursors with MTS was mainly related to their N-termini that define them as potential NatC substrates (F, L, l, W amino acid at position 2, green curve in Fig. 4e1) but that MTSs lacking this signature at position 2 contributed significantly to MTSs enrichment in the sel-TRAP data (blue curve in Fig. 4e1). This enrichment for proteins that are not putative NatC substrates was not observed for mitochondrial precursors lacking MTS (blue curve in Fig.4e2), as well as in the overall analysis of NatC sel-TRAP results (Fig. Sup3).

Finally, sel-TRAP analysis clearly demonstrated that mitochondrial proteins with an N-terminal MTS sequence are co-translationally recognized by NatC mainly when they have a leucine, a phenylalanine or to a lesser extent an isoleucine at position 2. Furthermore, unexpectedly, the observation that MTSs with other amino acids at position 2 were selectively enriched after NatC immunoprecipitation strongly suggests that other features of MTSs might be involved in their recognition by NatC.

### Substitution of leucine 2 by amino acids not overrepresented at position 2 of MTSs result in precursor accumulation and rescue of the dominant negative effect of an HSP60−13myc transgene

Finally, we explored the functional significance of selecting during evolution specific amino acids at the N-terminus of the mitochondrial addressing sequences. To this aim, we set up a mutagenesis experiment at position 2 of a typical MTS. We chose Hsp60p, an essential chaperonin of the mitochondrial matrix (35) as a model of the MTS-dependent import pathway because of (1) the presence of a highly conserved leucine at position 2 of its MTS (leucine 2 is observed in 13 of 17 Saccharomycotina yeasts studied) and (2) the availability of an antibody allowing sufficiently sensitive detection of its precursor when its mitochondrial import is impaired. In addition, to improve detection of Hsp60p precursors, we used the thermosensitive strain YPH499 pam16Δ-MAGN76D (36), which, even at the permissive temperature, exhibits a slight defect in mitochondrial import as evidenced by the detection of a low-intensity band corresponding to the precursor form of Hsp60p (Fig. 5a, Hsp60p WT lane). We used CRISPR/CAS9 technology to mutagenize this strain and create L2X genomic mutants of Hsp60p in which the leucine at position 2 was replaced by another amino acid (Fig. 5a, lanes corresponding to L2X mutants). Replacing leucine 2 in Hsp60p with aspartate, lysine, serine, valine or proline (Fig.5a L2D, L2K, L2S, L2V, L2P lanes) resulted in a dramatic increase in the intensity of the band corresponding to the Hsp60p precursor compared to that corresponding to its mature form, strongly suggesting an altered mitochondrial import of Hsp60p. In contrast, substitution of leucine with a phenylalanine, another hydrophobic amino acid over-represented at position 2 of the MTS-containing protein group, had no impact on the Hsp60p profile observed by western blot (Fig. 5a, L2P lane).

**Figure 5:**
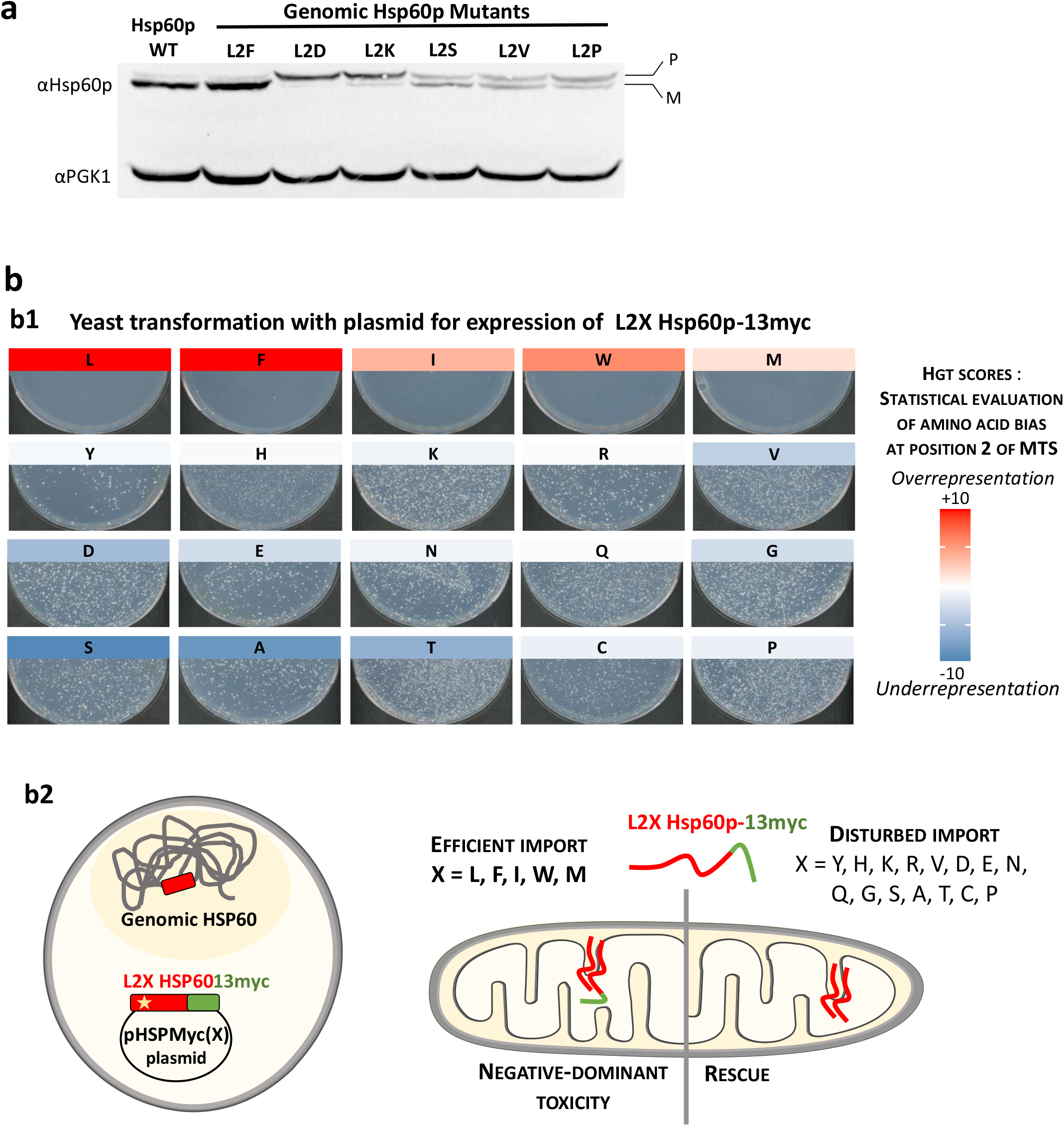
Mutagenesis of position 2 of Hsp60p MTS confirmed our predictions of the optimal residues for efficient mitochondrial import. (a) L2X genomic mutants of Hsp60p were generated using CRISPR/CAS9 technology (see Methods) and proteins were analyzed by SDS-PAGE followed by western blots using antibodies against Hsp60p and Pgk1p, used as loading control. The position of the precursor and mature Hsp60p are indicated. In L2D, L2K, L2S, L2V, L2P mutants, the increase in precursor bands relative to the mature band suggests a defect in mitochondrial import of the mutated Hsp60p protein. (b) The pHSPmyc(X) plasmids expressing different versions of Hsp60p−13myc mutated at position 2 were transformed into the thermosensitive strain YPH499 pam16Δ-MAGN76D (see b2 for experimental strategy). Photographs of the transformation plates (b1) showed that only a few clones were obtained after transformation with the plasmid encoding the wild-type leucine at position 2 of Hsp60p, a result probably due to the dominant-negative effect of the HSP60−13myc allele (see b2 for a schematic of the interpretation of results). Similar results were obtained after transformation of the strain with plasmids expressing the L2F, L2I, L2W and L2M mutants of Hsp60p−13myc. In contrast, transformations with plasmids expressing all other L2X mutants of Hsp60p−13myc resulted in more than 1000 transformants, which is likely due to impaired mitochondrial import of the mutated Hsp60p in these cases (see b2 for interpretation scheme). Interestingly, the two sets of residues at position 2 defined according to the results of the transformation experiment correlate perfectly with the amino acid usage biases at position 2 of the MTSs detected in our *in-silico* analysis, as indicated by the color scale representing the previously calculated HGT scores (Fig.4a1). The experimental strategy and interpretation scheme are presented in b2 (see main text for details).

In a second step, we developed a screen to easily test the effect of all possible amino acid substitutions at position 2 of Hsp60p. For this purpose, we took advantage of an HSP60−13myc allele that we found to have a dominant negative effect. Indeed, the transformation of *S. cerevisiae* with the pHSPmyc(L) plasmid, which allows the expression of the HSP60−13myc allele, led to no colony (plate L in Fig.5b1). To interpret these results, we hypothesized that the 13-myc C-terminal extension abolished the activity of Hsp60p C-terminal domain, previously shown to play a crucial role for the function of the protein (37, 38). Thus, since Hsp60p is essential for cell survival, we attributed the near absence of clones after transformation with pHSPmyc(L) to a dominant negative effect of the myc-tagged version of Hsp60 on the endogenous wild-type form. After being imported in mitochondria, Hsp60p forms a tetradecameric chaperonin complex in the mitochondrial matrix required for its own assembly (37). Thus, the association of the deficient protein Hsp60p-13myc with endogenous wild-type Hsp60p would lead to the inactivation of the Hsp60p complexes and the observed drastic loss of viability (Fig. 5b2). We further observed that an altered mitochondrial import of Hsp60p, as induced by the cis mutations L2D, L2K, L2S, L2V, L2P (Fig. 5a), resolves the dominant negative effect of the 13-myc tag. More than 1,000 transformants were obtained with the pHSPmyc(X) plasmids corresponding to these mutations, where X represents the amino acid that replaces leucine at position 2 of Hsp60 in the Hsp60p-13myc protein (Fig. 5b1).

Strikingly, after testing all possible amino acids at position 2 of Hsp60p-13myc, we found that the two groups of amino acids defined on the basis of their ability to rescue C-terminal tag toxicity correlated strongly with their enrichment at position 2 of *S. cerevisiae* mitochondrial addressing sequences, as assessed by our statistical analyses (Fig. 2 and Fig.5b). Indeed, the amino acids that did not result in more than 1−3 clones after transformation were those we showed to be to be enriched at position 2 of MTSs, i.e., leucine (HGT=71), phenylalanine (HGT=15), isoleucine (HGT=4), tryptophan (HGT=6), and methionine (HGT=1.6). Western blot analysis revealed that the few clones obtained for these 4 amino acid substitutions did not express Hsp60p-13myc protein and likely escaped toxicity through plasmid mutations, which was confirmed by sequencing some of these plasmids (Fig. Sup4b). For all other amino acids (except tyrosine which showed a lower number of transformants), we observed more than 1,000 colonies after transformation by pHSPmyc(X) plasmids. Western blot analysis of the resulting clones confirmed that they all express Hsp60p-13myc protein at a level roughly comparable to that of the endogenous protein (Fig. Sup. 4a). The only exceptions are the clones transformed with pHSPmyc(Y) in which the level of Hsp60−13myc protein was systematically reduced (Fig. Sup4b). In addition, we consistently observed, in all clones with a rescue phenotype, an Hsp60p-myc band corresponding predominantly to the non-imported precursor form (see for example the profile of the L2Q mutant for which both bands are visualized with the anti-myc antibody) (Fig. Sup4a). More surprisingly, we also observed a defect in import and/or processing of the endogenous, wild type, Hsp60p, reflected by the clearly visible band corresponding to the precursor form of the protein (Fig. Sup4a). This observation may indicate that a small fraction of the Hsp60p-13myc proteins may have been imported, resulting in a significant, although not lethal, loss of Hsp60p function that could be associated with impaired processing of mitochondria-targeted endogenous Hsp60p precursors, as previously observed upon inactivation of mitochondrial chaperonin (35, 39).

The results obtained in our mutagenesis experiments allowed us to demonstrate the dramatic functional consequences of modifying the residue located at position 2 of the mitochondrial addressing sequence. These results, which fully validated our *in-silico* predictions, also supported the existence of strong evolutionary constraints that shaped the current amino acid biases at position 2 of MTSs, thus ensuring an optimized mitochondrial import.

## DISCUSSION

In this work, we developed an *in-silico* approach to identify functional groups of proteins with N-terminal amino acid usage biases. The resulting functional mapping of N-terminal residues reveals possible evolutionary constraints at position 2 of nascent chains and highlights potential early mechanisms of protein regulation or targeting. In particular, we discovered the potential critical role of the residue located at position 2 of mitochondrial N-terminal targeting sequences in *S. cerevisiae*. This prediction was experimentally validated using an original system based on a dominant-negative allele of HSP60, that we set up to test the efficiency of mitochondrial import after mutagenesis at position 2 of the MTS. Strikingly, we observed that only the 5 hydrophobic amino acids (Leu, Phe, Ile, Trp and Met) over-represented at this position allowed efficient mitochondrial import of Hsp60p protein. This finding raises the question of the nature of selective pressures acting specifically on position 2 of the MTSs in the context of their other well-established characteristics. It also strongly suggests the involvement in optimizing mitochondrial precursor import of N-terminal modification enzymes, including NatC, which we found to co-translationally interact with MTS-bearing precursors.

### The N-terminal signature of MTSs in light of the current knowledge about mitochondrial import mechanisms

MTSs are typically 15−50 amino acid residues in length and form positively charged amphipathic α-helices. It has been suggested that these sequences possess several biochemical features involved in their interaction with binding sites of the mitochondrial translocation machinery (26). First, the amphipathic alpha helix interacts with two outer membrane translocase (TOM) receptors: the hydrophobic surface is recognized by Tom20 and the positively charged one by Tom22 (40–42). In addition, translocation of the precursor into mitochondria can occur via an acidic binding chain involving the interactions between MTSs basic residues and some residues from Tom7p, Tom22p, and Tim23p (43). Finally, the mitochondrial membrane potential (Δψ) generates an electrophoretic force on the MTSs and the net positive charge of the MTSs is the critical determinant of this effect (44, 45). Taken together, these data indicate that the hydrophobicity of MTSs is mainly determinant for their primary interaction with the outer mitochondrial membrane receptor Tom20p.

In addition, TargetP2.0, a new N-terminal sorting signal detection tool based on a machine learning approach, has recently further improved the prediction of mitochondrial presequences (46). By analyzing the attention layer of their final network, these authors found that the residue at position 2 of the presequences had a strong influence on their classification. More specifically, they observed that, in fungi, peptides targeted to the endoplasmic reticulum (ER) and to mitochondria showed at position 2 an underrepresentation of residues that allow the removal of the initiator methionine, i.e. Ala, Cys, Gly, Ser, Thr, or Val (46). This observation is consistent with the results we obtained in *S. cerevisiae* (Fig2b). Indeed, we found that only 20% of MTSs have amino acids at position 2 allowing the removal of their iMet compared to 56% within the proteome and 69% among cytosolic proteins. Of note, Forte et al. also previously observed that less than 25% of ER signal sequences would be predicted to be MetAPs substrates and further demonstrated that removal of iMet from carboxypeptidase Y precursor inhibits its ER translocation (47). Similarly, we can speculate based on our results and on the observation made by Almagro Armenteros et al. that removal of iMet may be deleterious to mitochondrial import.

In favor of this hypothesis, we observed that the optimal residues at position 2 of MTSs, i.e. Leu, Phe, Ile, Trp and Met, have high hydrophobicity indices, but also that none of them allow iMet removal. Conversely, the small residues Ala and Cys, which are also highly hydrophobic but induce iMet cleavage when positioned at position 2 of the nascent chains, are underrepresented at this position of the yeast MTSs. Therefore, the N-terminus of most MTSs will be characterized by the sequence M(L/F/I/W/M) with two consecutive hydrophobic residues in positions 1 and 2 and this hydrophobic N terminus may be important in the early binding of mitochondrial precursors to the mitochondrial surface. Indeed, several analyses indicate that consecutive hydrophobic residues could be involved in the binding of MTSs to Tom20p. In particular, by measuring the binding of a peptide library to Tom20, a consensus sequence θφχβφφ was identified (where θ, β, φ, and χ represent a hydrophilic, basic, hydrophobic, and any residue, respectively) and proposed as a Tom20 binding motif (48). Bioinformatic analysis extended this consensus by identifying statistically significantly enriched hexamers in the 30 first residues of MTSs, and their detection was implemented in MitoFates, one of the most efficient tools for MTSs prediction to date (49). Most of these hexamers showed strong potential to form amphiphilic helices and includes several consecutive hydrophobic residues.

By integrating these data from the literature and our results on MTS position 2, we propose a model explaining the critical role of hydrophobic residues at the N-terminus of mitochondrial addressing sequences (Fig. 6). We can hypothesize that the selection of large hydrophobic amino acids at position 2 of MTSs was a successful evolutionary solution, the consequences of which were high N-terminal hydrophobicity due to (1) conservation of the hydrophobic iMet, and (2) possible N-terminal neutralization of the N-terminal amine by acetylation by NatC. Positioning the hydrophobic site directly at the N-terminus of MTS would then allow for early interaction with Tom20p and the mitochondrial membrane. Consequently, the arginines that provide the positive charge necessary for translocation of the precursors to mitochondria could be highly enriched from residue 3 of MTS, without compromising the interaction with Tom20p.

**Figure 6:**
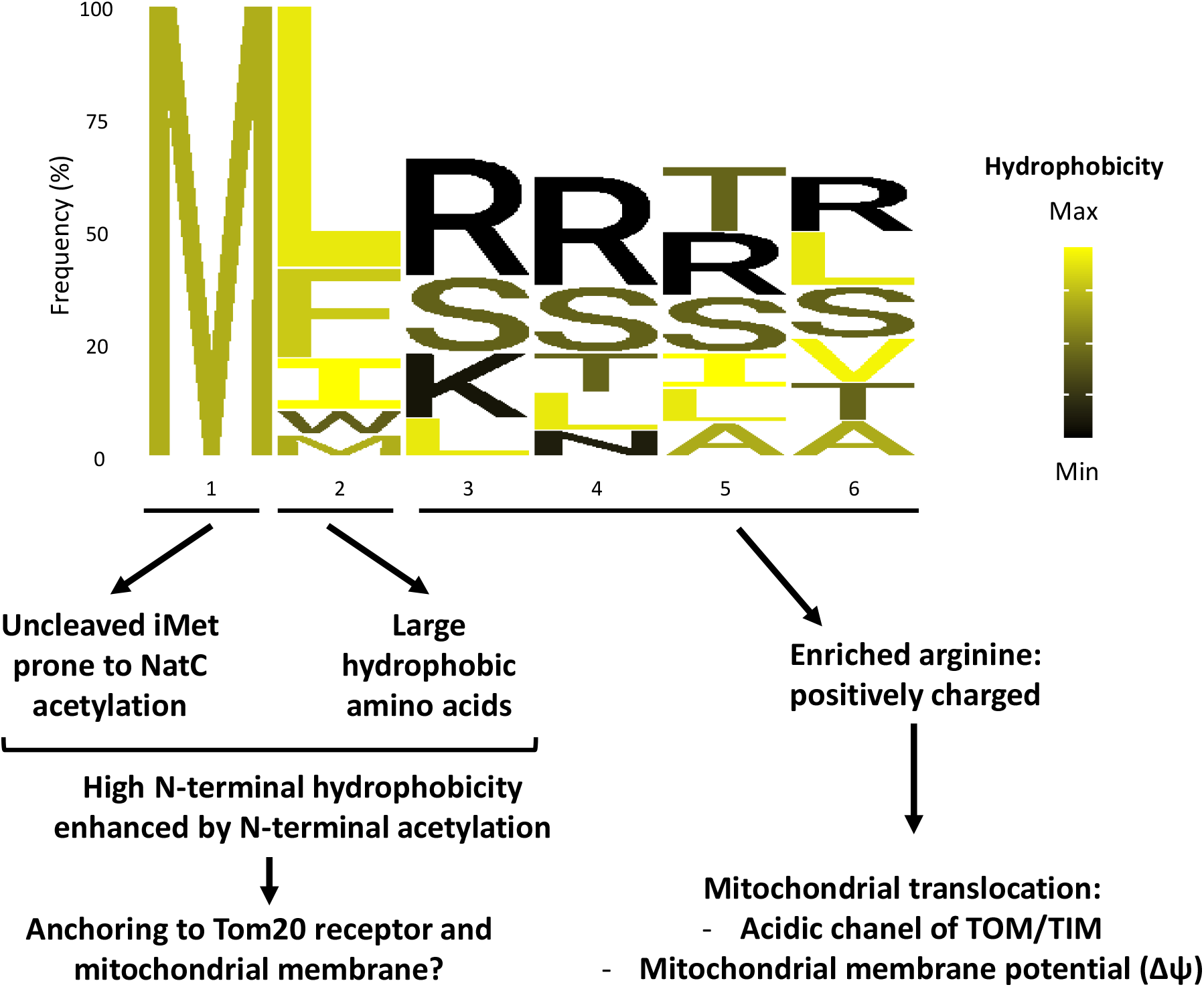
A model to explain the bias at position 2 of MTSs and the importance of their N-terminal hydrophobicity. Sequence logo (77) of the first 6 residues of *S. cerevisiae* MTSs with optimal residues at position 2 (Leu, Phe, Ile, Trp, Met) according to our amino acid usage bias analysis and our results of Hsp60p MTS mutagenesis. For residues 3 to 6, only the most frequent residues that represent at least 60% of the observed residues are represented. The N-terminus of this class of MTS exhibits high hydrophobicity due to (1) the lack of iMet cleavage, (2) the intrinsic hydrophobicity of the residue at position 2, and (3) potential acetylation of the N-terminal amine by NatC. We propose that this N-terminal feature is favorable for an efficient interaction of MTSs with the mitochondrial membrane and the Tom20p receptor. In addition, we observe, from the residue at position 3 of MTSs, a high enrichment of positively charged arginine involved in the translocation of MTSs into mitochondria. We propose that this residue configuration in MTSs has been selected during evolution because of its ability to promote efficient mitochondrial import with limited interference between the two driving forces, hydrophobicity vs. positive charge, required in this mechanism.

### A conserved role of the N-terminal signature in the mitochondrial import of MTS-bearing proteins

In support to our observation of significant biases at position 2 of the MTS in yeast, our mutagenesis experiments using Hsp60p as a model revealed the importance for proper mitochondrial import of having at this position large hydrophobic amino acids that do not lead to cleavage of the initiator methionine. Only leucine, phenylalanine, isoleucine, tryptophan and methionine at position 2 did not affect the mitochondrial import of our dominant negative Hsp60p-13myc allele.

Strikingly, mutagenesis of position 2 of the rat aldehyde dehydrogenase (Aldh) MTS, whose first residues are MLRAAL, yielded similar results to those we obtained by mutating Hsp60p in yeast (50). Replacement of leucine at position 2 with a valine, which is also a hydrophobic residue but which drives the removal of iMet, did not allow efficient *in vitro* import of Aldh, whereas isoleucine, tryptophan, or phenylalanine at position 2 had no impact on mitochondrial import. Interestingly, the correct import of the L2V mutant was restored when it was translated using a system that did not remove the hydrophobic iMet. Similarly, a mutation that increased the N-terminal hydrophobic surface area compensated for the L2V mutation. These data underscore the general importance during eukaryotic evolution of the N-terminal hydrophobicity of MTSs and the necessary balance with positive charges that are also critical for precursor translocation into mitochondria. In this context, it would be crucial to further study the evolution of MTSs in order to better understand the characteristics that allow their optimized targeting to mitochondria.

### The characteristics of MTSs allow their efficient recognition by the N-terminal acetyltransferase NatC

In this work, we discovered the specific and optimal N-terminal signature of mitochondrial targeting sequences, defined by the presence of a hydrophobic amino acid directly after the uncleaved initiator methionine. This led us to hypothesize that, in *S. cerevisiae*, mitochondrial precursors with a MTS could be cotranslationally targeted by NatC.

Interestingly, previous N-terminal acetylome analyses in *S. cerevisiae* detected three mitochondrial precursors: two of them had an ML-terminus, Pda1p and Glr1p, and were fully acetylated (51). However, no global experimental identification of NatC targets in yeast had been achieved before our work. Therefore, we used the Selective Affinity Purification of Translating Ribosomes (sel-TRAP) technology to identify nascent chains co-immunoprecipitated with NatC. In this experiment, 30% of the captured nascent chains identified by their corresponding mRNA were mitochondrial precursors with a MTS, most of them having a leucine or phenylalanine at position 2. However, the nature of the residue at position 2 may not be the sole determinant of the affinity of NatC for these precursors.

The structure of NatC bound to a *S. cerevisiae* ligand highlighted the importance of residue at position 3 of the substrates that is fully involved in the binding in the catalytic pocket. More specifically, the analysis of the ligand bound structure revealed that when an arginine is present at position 3 of the peptide, it binds to a specific site in the catalytic pocket and strongly increases the catalytic efficiency of NatC towards the peptide measured in a catalytic assay (52). Hence, many mitochondrial precursors have features for optimal NatC binding (retention of iMet, hydrophobic residue at position 2 and arginine at position 3), as depicted in figure 6. As a matter of fact, one-third of MTSs that have a leucine at position 2 also have an arginine at position 3.

In summary, it appears that many MTSs possess N-terminal features that are optimal for efficient binding and acetylation by NatC. Using sel-TRAP technology, we have indeed demonstrated that MTSs are recognized co-translationally by NatC, suggesting their early N-terminal acetylation. More surprisingly, we also observed that some MTSs without features compatible with binding to the active site of NatC, i.e., those for which the i-Met is cleaved, were enriched in our sel-TRAP data. It is then tempting to speculate that other features of these MTSs, besides their N-terminal composition, could allow their recognition by NatC.

### A role for NatC in mitochondrial biogenesis?

From yeast to human, phenotypic analyses of cells lacking NatC have highlighted its critical importance for mitochondrial function. In *S. cerevisiae,* deletion of any of the NatC subunits has been shown to induce growth defects on non-fermentable carbon sources, such as glycerol and ethanol (52–56).

The importance of NatC for mitochondrial function was also underscored by the observation that depletion of NatC catalytic subunit, Naa30, in a human cell line resulted in reduced levels of mature mitochondrial matrix membrane proteins, many of whose precursors were substrates of NatC (57). This phenotype was associated with loss of mitochondrial membrane potential and fragmentation of mitochondria.

Furthermore, in glioblastoma-initiating cells (GICs), that drive tumorigenesis of the most common primary brain malignancy, a strong overexpression of the protein Naa30 was detected. shRNA-mediated knockdown of *NAA30* gene in GICs demonstrated that NatC may serve as a therapeutic target in these types of cancer, as intracranial transplantation of such NatC depleted cells resulted in prolonged animal survival compared to control mice transplanted with unmodified GICs (58). At the cellular level *NAA30* knockdown in GICs cells results in reduced viability and decreased mitochondrial hypoxia tolerance, with a more abrupt and severe mitochondrial membrane depolarization compared to control cells in such condition. These observations suggested that NatC contribute to the stability of the mitochondrial potential, which was supported by the observation that several genes involved in hypoxia response, including HIF1α, a central regulator of transcriptional response to hypoxia, were upregulated in absence of NatC even in normal culture conditions.

We have shown that, in the yeast *S. cerevisiae*, a large fraction of proteins harboring MTSs is susceptible to acetylation by NatC during their translation. At the molecular level, N-terminal acetylation by NatC was shown to modulate proteins ability to interact with other proteins or membranes allowing protein complexes formation or targeting to specific subcellular subsites. Examples of such NatC-dependent interactions include the formation of the Gag capsid of the yeast L-A helper virus (28, 59), or of the Ubc12-Dcn1 complex (60, 61), the insertion of the tRNA-specific methyltransferase Trm1-II into the inner nuclear membrane (62), the targeting of the human ADP-ribosylation factor-like protein 8b to lysosomes (63), or the localization of Arf-related protein 1 (hARFRP1) and its yeast orthologue yArl3 to the trans-Golgi via an interaction of their acetylated N-terminus with the hSys1/Sys1p protein (64, 65).

Similarly, it is tempting to speculate that N-terminal acetylation of MTSs further increases their N-terminal hydrophobicity, which may enhance their binding to the mitochondrial receptor Tom20p and their subsequent mitochondrial addressing. A previous study of the GFP localization pattern of potential NatC substrates, including three mitochondrial proteins, failed to identify altered localization in yeast lacking the catalytic subunit of NatC (51). However, the fluorescent imaging approach used in this study may not be sensitive enough to detect subtle mitochondrial import defects that could be induced by the loss of N-terminal acetylation of MTSs in the absence of NatC. Moreover, N-terminal acetylation could have various effects on mitochondrial addressing efficiency, depending on other characteristics of the MTS, including its overall N-terminal hydrophobicity. Thus, the mitochondrial phenotype of cells lacking NatC activity could be due either to the cumulative effect of mild defects in the import of many mitochondrial proteins or to strong effects on a limited number of mitochondrial precursors for which N-terminal acetylation by NatC would be critical. A systematic exploration of the impact of NatC depletion on the various MTS-bearing precursors is still needed to link the strong mitochondrial phenotypes of NatC-depleted cells to the specific role of N-terminal MTS acetylation on mitochondrial precursors fate.

## MATERIAL AND METHODS

### S. cerevisiae strains, plasmids, and growth conditions

All *Saccharomyces cerevisiae* strains used in this study are described in Supplementary Methods (Table S5). The plasmids used for yeast transformation are also described in the Supplementary Methods (Table S6) as well as the oligonucleotides used for the diverse genetic constructs (Table S7 and S8).

The strains used for the sel-TRAP experiment were derived from BY4741 by homologous recombination with the ProtA-His5 cassette amplified from the pBXA plasmid (66). The temperature-sensitive and respiratory-deficient strain YPH499 pam16Δ-MAGN76D was kindly provided by Agnes Delahodde with the corresponding reference strain YPH499 (36). The substitutions of the leucine at position 2 of Hsp60p were obtained by site-directed mutagenesis using CRISPR/CAS9 technology. YPH499 strains were co-transformed with the plasmid pAEF-HSP60 (67) encoding the Cas9 endonuclease and expressing a guide RNA targeting the HSP60 sequence, and a HSP60 repair template including the desired mutation. The resulting strains were used for PCR amplification of the mutant HSP60 sequences to obtain the diverse pHSPmyc(X) plasmids (X representing the amino acid at position 2 of Hsp60p sequence) by cloning into pZMYA7 (32), a pRS416-derived plasmid allowing C-terminal tagging with the 13Myc epitope.

Yeast were grown on rich medium (1% bactopeptone, 1% yeast extract). When required for plasmid selection, cells were grown on CSM medium (0.17% yeast nitrogen base, 0.5% ammonium sulfate, 2% dextrose, 0.07% CSM mixture) depleted of the selective compound. *Hygromycin* B (200 μg/ml) was used in rich medium to select mutants containing pAEF5-derived plasmid after CRISPR/CAS9 procedure. After genome sequencing verification, plasmid was removed from HSP60 mutant strains by overnight culture in rich medium without antibiotic selection.

### Antibodies

The following antibodies were used for western blotting or immunoprecipitation: rabbit IgG-HRP polyclonal antibody (PAP; code Z0113; Dako), rabbit Hsp60 polyclonal antibody (gift from J.P. Di Rago), anti-PGK1 mouse monoclonal antibody (ab113687, abcam) and mousse moclonal anti-myc antibody (11667149001, Roche).

### Data for the *in-silico* analyses

The 6,050 sequences of *S. cerevisiae* proteome (proteome ID: UP000002311) were downloaded from the UniProtKB database (68) (http://www.uniprot.org/uniprot/?query=proteome:up000002311&format=fasta). Since proteins encoded by transposons have redundant sequences and their distribution vary depending on yeast strains (69), the 91 proteins annotated as transposons according to the Saccharomyces Genome Database (https://www.yeastgenome.org/) were removed from the proteome.

Gene Ontology annotations were downloaded from the GO consortium database (70, 71) (http://current.geneontology.org/annotations/sgd.gaf.gz). For extensive analysis of mitochondrial proteins, the 1,190 proteins belonging to the mitochondrion Gene Ontology term (GO:0005739) were manually curated: proteins that were associated to mitochondria based on only one high-throughput experiment were removed, leaving 726 proteins for which association with mitochondria was strongly evidenced. Curated mitochondrial proteins were then sorted depending on whether they carry an N-terminal mitochondrial targeting sequence, according to sequence annotations provided by the UniProtKB database (68). 361 over 726 mitochondrial proteins were annotated as carrying a MTS (Table S2).

For the comparative analysis, 17 species were selected from the Saccharomycotina yeast lineage (23) (see Table S9 for the list of selected species). Orthology groups recently established for 322 budding yeast species (24) were used to retrieve the 311 groups containing a mitochondrial protein from *S. cerevisiae*, among which 161 had an N-terminal MTS. More than 95% of these groups (298) contained orthologs for at least 13 of the 17 selected species, and all recovered orthologs for at least 9 of these species.

### HGT scores derived from hypergeometric test

Hypergeometric tests (HGT) have been classically used to identify significant differences between the observed frequencies (F_obs_) in various subsets of interest and the reference frequency (F_ref_) of the corresponding input set. The p-values provided by these tests were then used to calculate hypergeometric scores (HGT scores) as follows: |HGT score|= log10(P-value), with a negative sign assigned to the score in case of underrepresentation (F_obs_< F_ref_).

### Analysis of amino acids usage bias at position 2 in the proteome

For each of the 20 amino acids, the frequency of use at position 2 (F2) was calculated and compared with the respective average use of the amino acid in the proteome (Fmean). Differences were statistically evaluated using hypergeometric tests and HGT scores were calculated. The observed frequency value at position 2 was also used to define a reference value called F2ref calculated as follows: if F2>Fmean, F2ref= 0.9*F2, otherwise F2ref= 1.1*F2. The F2ref value was then used to test the observed frequencies at any other position and calculate an aspecificity score, representing the percentage of positions with a frequency greater than F2ref when F2>Fref, or less than or equal to F2ref when F2≤Fref. To calculate this score, the first 500 positions of the proteome were analyzed by a 100-position long sliding window. At each step of the analysis, the window slides by one position and the number of positions within the window with a frequency value higher (when F2>Fref) or lower (when F2≤Fref) than the reference value F2ref was evaluated, to calculate a local aspecificity score. The final retained aspecificity score corresponds to the maximum value of the measured local aspecificity scores.

### Analysis of the amino acid usage bias at position 2 in subsets of proteins of the Gene Ontology

We have implemented a set of functions in R to detect gene ontology terms showing, among their associated proteins, a significant and specific amino acid usage bias at position 2. The gene annotations were downloaded from the GO consortium and stored in MGSA objects (R package MGSA (72)). The second residue associated with each protein was extracted from the downloaded proteome (UniProtKB database). For each GO domain (cellular component, molecular function, biological process), contingency tables were generated that count, for all GO terms, the number of proteins with a given amino acid at position 2. To avoid detecting amino acid bias due to proteins with the same N-terminal sequence, proteins sharing the same first 10 residues were grouped under the same identifier, which was counted only once. These contingency tables were used to calculate HGT scores and detect GO terms with significant over-representation relative to the proteome (HGT score>4) of one or more amino acids in position 2 of their associated proteins. GO terms that showed the lower over-representation (fold change <1.8) and mainly correspond to GO term regrouping very large number of proteins were first removed from this initial list. Then, to reduce the redundancy of the detected GO terms, we developed a two-step algorithm to (1) select the minimum number of low-redundant GO terms (overlap of GO terms with larger ones <40%) maximizing the number of proteins associated with the detected amino acid usage biases (BestN GO), and (2) identify smaller low-redundant GO terms (overlap with larger GO term<30%) that are partially or fully included in one BestN GO (overlap≥40%) terms but have at least 30% higher frequency bias (BestF GO). Finally, to verify that the amino acid usage bias detected in a given GO term was not due to a specific N-terminal composition of the associated proteins, the search for position-specific amino acid bias was extended by calculating the HGT enrichment scores relative to the proteome on the first 100 positions of these proteins. Details of the GO reduction algorithm are available in the supplementary methods.

### Selective Translating Ribosome Affinity purification (sel-TRAP)

For analysis of canonical ribosomes and NatA, NatB and NatC-associated ribosomes, ribonucleoparticle purifications from cells expressing proteinA-tagged baits (respectively Rpl16a, Ard1, Nat3 and Mak3) were performed essentially as described in (32). Briefly, frozen grindates obtained from cells grown in rich medium were homogenized in nine volumes of extraction buffer (20 mM Hepes pH 7.5, 110 mM KOAc, 2 mM MgCl_2_, 0.1% Tween-20, 0,5% Triton X−100, 1 mM DTT, 1× protease inhibitors cocktail, complete EDTA-free, Roche and antifoam B, Sigma, 1:5000). The obtained extract was clarified by filtration through 1.6 μm GD/X Glass Microfiber syringe filters (25 mm, Whatman) and input samples were collected for analysis of total cellular RNA and protein content. The immunoprecipitation of proteinA-tagged baits was performed by further incubation of the extract for 30 min at 4°C with IgG-conjugated magnetic beads. Beads were then washed three times with extraction buffer, once with washing buffer (0.1 M NH4OAc, 0.1 mM MgCl_2_) supplemented with 0.02% Tween-20 and four times with washing buffer without Tween−20. Beads were then split into two samples for protein and RNA analysis respectively. Immunoprecipitated proteins were eluted with 0.5 M NH_4_OH, 0.5 mM EDTA, lyophilized and resuspended either in SDS-sample buffer for SDS-PAGE or in 25 mM ammonium carbonate for mass spectrometry analysis. Immunoprecipitated RNA was eluted by using the RLT buffer from Quiagen RNeasy Mini Kit@ that was used to purified total and immunoprecipitated RNAs.

### Mass spectrometry analysis of sel-TRAP experiment

Mass spectrometry analysis procedures, performed at Proteomics Core Facility of Institut Jacques Monod, are described in Supplementary methods, and mass spectrometry proteomic data were deposited on the ProteomeXchange Consortium (73) via the PRIDE partner repository with dataset identifier PXD034922.

Mascot score distributions of immunoprecipitations from control (unlabeled BY4741) or protein A-labeled strains were compared to distinguish specifically interacting proteins from background. Based on the distribution of Mascot scores in the control, a cutoff score of 250 was chosen to retain less than 5% false positives. Only proteins with a Mascot score greater than 250 in the immunoprecipitation data of protein A-labeled strains were considered for further analysis. Proteins with a score greater than 250 in the control or with a ratio of protein A-labeled strain to control less than 2 were excluded in the final list of partners detected in sel-TRAP experiments.

Venn diagrams obtained from two independent experimental sets were used to compare the protein partners of Ard1, Nat3 and Mak3 and determine the shared and specific partners observed in a reproducible manner. In this representation, for a more accurate definition of partners, all proteins with at least a 5-fold higher Mascot score in one of the sel-TRAP experiments compared with the others was assigned as specific partner of the corresponding immunoprecipitated protein.

### Microrray analysis of sel-TRAP experiment

The microarray data and the related protocols are available at ArrayExpress website (https://www.ebi.ac.uk/arrayexpress/experiments/E-MTAB-11772/) with the dataset identifiers E-MTAB-11772.

After sel-TRAP experiment, input and immunoprecipitated (IP) RNAs were purified using the RNeasy Mini Kit (Qiagen). One μg of RNA from each sample was then reverse-transcribed and labeled with Cy3 or Cy5 dye using the indirect labeling procedure. cDNAs of input and IP samples were then hybridized on Agilent’s ab 8×60K *S. cerevisiae* custom DNA chip (AMADID: 027945). Arrays were read using an Agilent scanner at 2 μm resolution and the signal segmentation was done using the feature extraction software (Agilent). The data was normalized without background subtraction using the global Lowess method (74).

To access the specifically enriched mRNAs in the Ard1, Nat3, or Mak3 sel-TRAP experiments, the measured log2(IP/input) ratios were corrected by subtracting the corresponding value obtained from the Rpl16a immunoprecipitation. Average specific enrichment values were then calculated from two independent experiments. The obtained data were filtered according to the mRNA enrichment values using a threshold value increasing in 0.1 steps from −3 to +4. From the lists of mRNAs detected above these thresholds, the evolution of the proportions of the 20 amino acids at position 2 of the corresponding proteins was analyzed to identify any possible enrichment related to this position. HGT scores were also calculated for each threshold value to validate the statistical significance of the detected enrichments. After this analysis, the maxima of the HGT scores were observed for an enrichment threshold value around 0.8 (see Figure 4a) and this value was then used for subsequent analyses of Mak3-associated mRNAs. The LIMMA algorithm (75, 76) statistically validated the reproducibility of the list of enriched mRNA obtained with this threshold: all mRNAs with a mean log2 ratio greater than 0.8 in the two independent sel-TRAP experiments had an adjusted p-value lower than 0.05.

## ACKNOWLEDGEMENT

We are grateful to Jean-Paul Di Rago and Deborah Tribouillard for the fruitful discussions on mitochondria and suggestions of experimental systems. Our thoughts go to the late Agnes Delahodde who provided us with the heat-sensitive yeast strain that proved crucial for the validation of our *in-silico* prediction.

## FUNDING

This research was funded by the Emergence program of the Sorbonne University, the Systems Biology Network of the Paris Seine Biology Institute, and Fondation ARC pour la recherche sur le Cancer, https://www.fondation-arc.org/recherche-cancer (PJA 20171206624). The PhD of El Barbry H. was funded by the doctoral school Complexité Du Vivant of Sorbonne Université, and the PhD of Nashed S. by the program for disabled students of the French National Center for Scientific Research.

## CONFLICT OF INTEREST

The authors declare that they have no competing interests.

## SUPPLEMENTARY INFORMATION

### TABLE OF CONTENT

The supplementary information refers to additional data on methods and results, and includes several tables.

Some of the additional results are available as supplementary figures S1 to S4.

Tables S1-S4 are available as independent excel files. Table S5-S9 are grouped in Supplemental Methods.

**Table S1: Complete table of GO terms identified with amino acids biases at position 2**

**Table S2: Annotation of mitochondrial proteins of S. cerevisiae**

**Table S3: Microarray data from sel-TRAP experiments**

**Table S4: Mass spectrometry data from sel-TRAP experiments**

**Table S5: Yeast strains**

**Table S6: Plasmids and derived strains**

**Table S7: Oligonucleotides**

**Table S8: Codon used in the reparation cassette to encode the desired X mutation.**

**Table S9: List of the yeast species used for the genomic comparative studies**

**Fig Sup 1:**
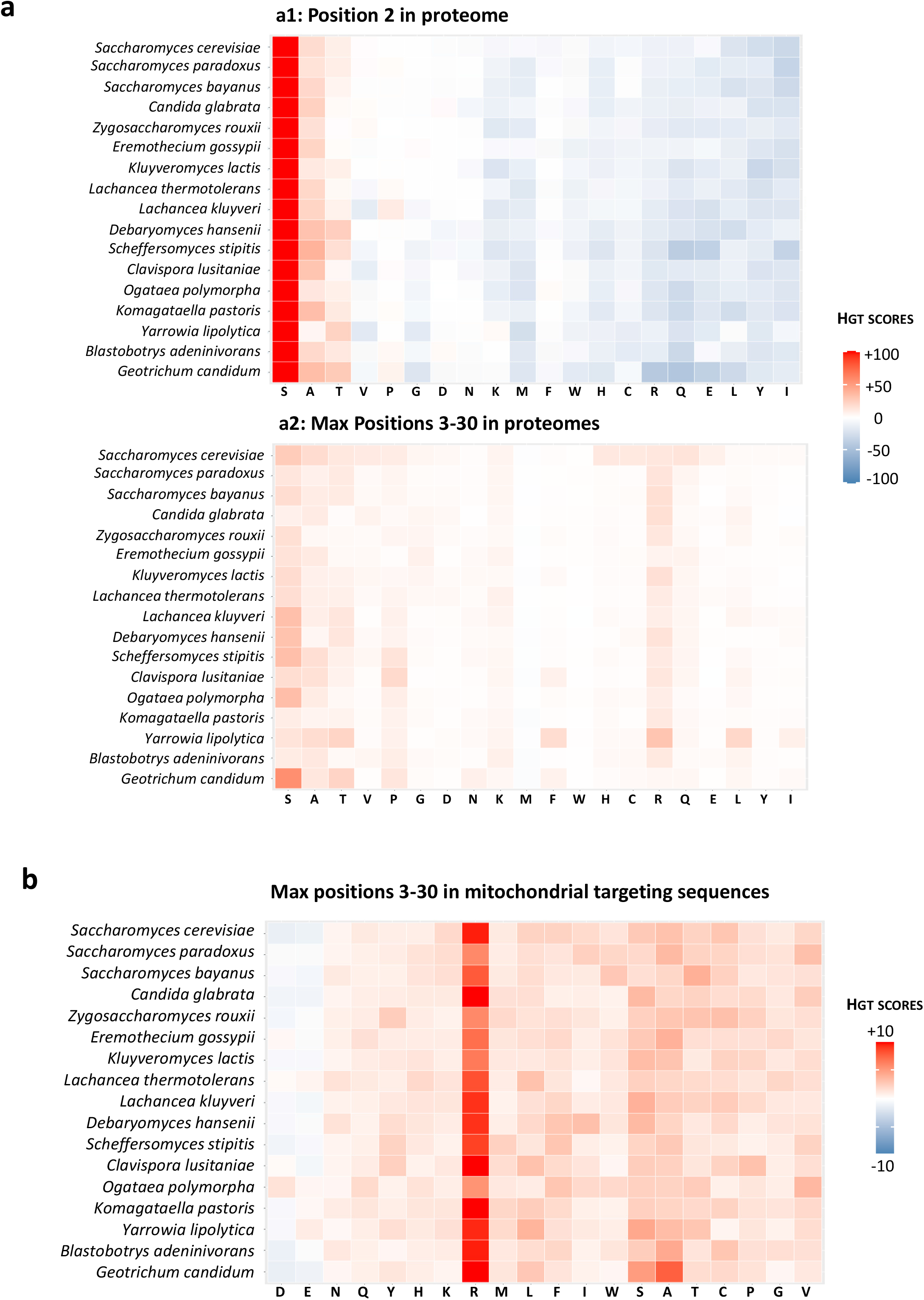
Conservation and specificity analysis of positional bias in amino acid utilization in the Saccharomycotina lineage. (a) Heatmap showing, for 17 budding yeasts of the Saccharomycotina lineage, the hypergeometric enrichment scores (HGT scores) obtained by comparing the utilization of amino acids at position 2 of the proteome to their respective average usage at any position (a1). Amino acid biases at position 2, particularly the high enrichment of serine, are similar in all species analyzed. The maximum values of the HGT scores obtained by comparing the amino acid usage at each position between 3 and 30 in the proteome to the average usage (a2) do not reach the values observed at position 2, as observed in the heatmap presented in a2 which confirmed the specificity of the amino acid bias observed at position 2. (b) Heatmap showing, for 17 budding yeasts of the Saccharomycotina lineage, the maximum values of the HGT scores obtained by comparing the amino acid utilization at each position between 3 and 30 of the MTSs with the amino acid usage in the proteome at these same positions. Comparison with the heatmap analyzing the amino acid bias at position 2 of the MTSs (Fig. 2c) clearly validated the specificity of the biases detected at position 2 of MTSs (i.e. enrichment in L, F and to a lesser extend I and W).

**Fig Sup 2:**
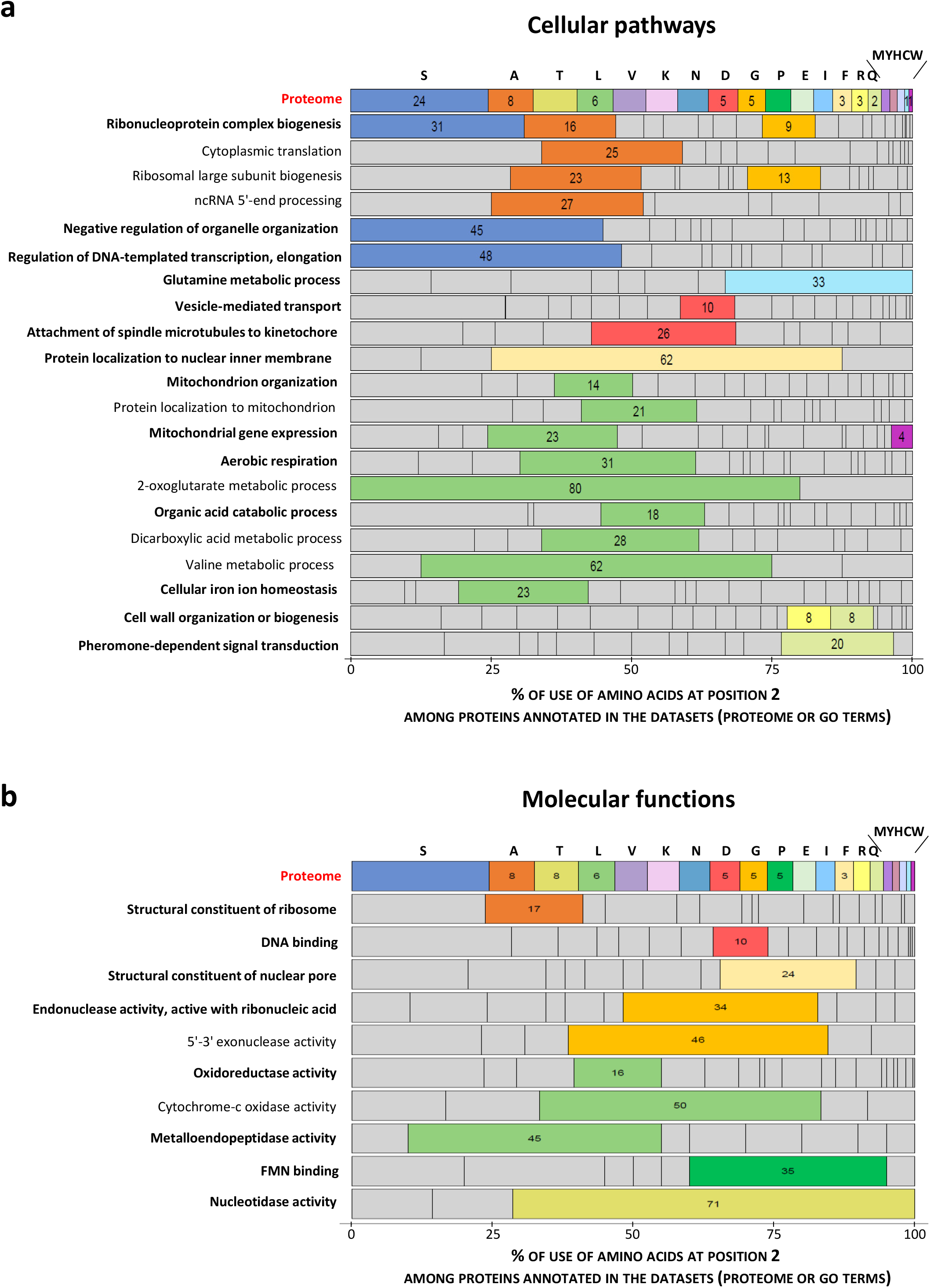
Amino acid usage biases at position 2 in GO terms related to “Cellular pathways” and “Molecular functions” GO categories. Barplot showing the percentages of use of the 20 amino acids at position 2 in retained GO terms related to cellular pathways (a) and molecular function (b) compared to their use at the same position in the proteome. bestN GO terms corresponding to the minimal number of terms maximizing the number of proteins with the detected usage biases (see Methods) are indicated in bold. For a better characterization of position 2 functional biases, bestF GO terms were retrieved also by our algorithm (regular typography). Each of them contains proteins included in one BestN GO term and has higher position 2 frequency biases. For each GO term, highlighted amino acids are those for which the percentage of use at position 2 is significantly higher than that observed at the same position in the proteome (HGT score > 3). Amino acids are sorted in decreasing order of use at position 2 in the

**Fig Sup 3:**
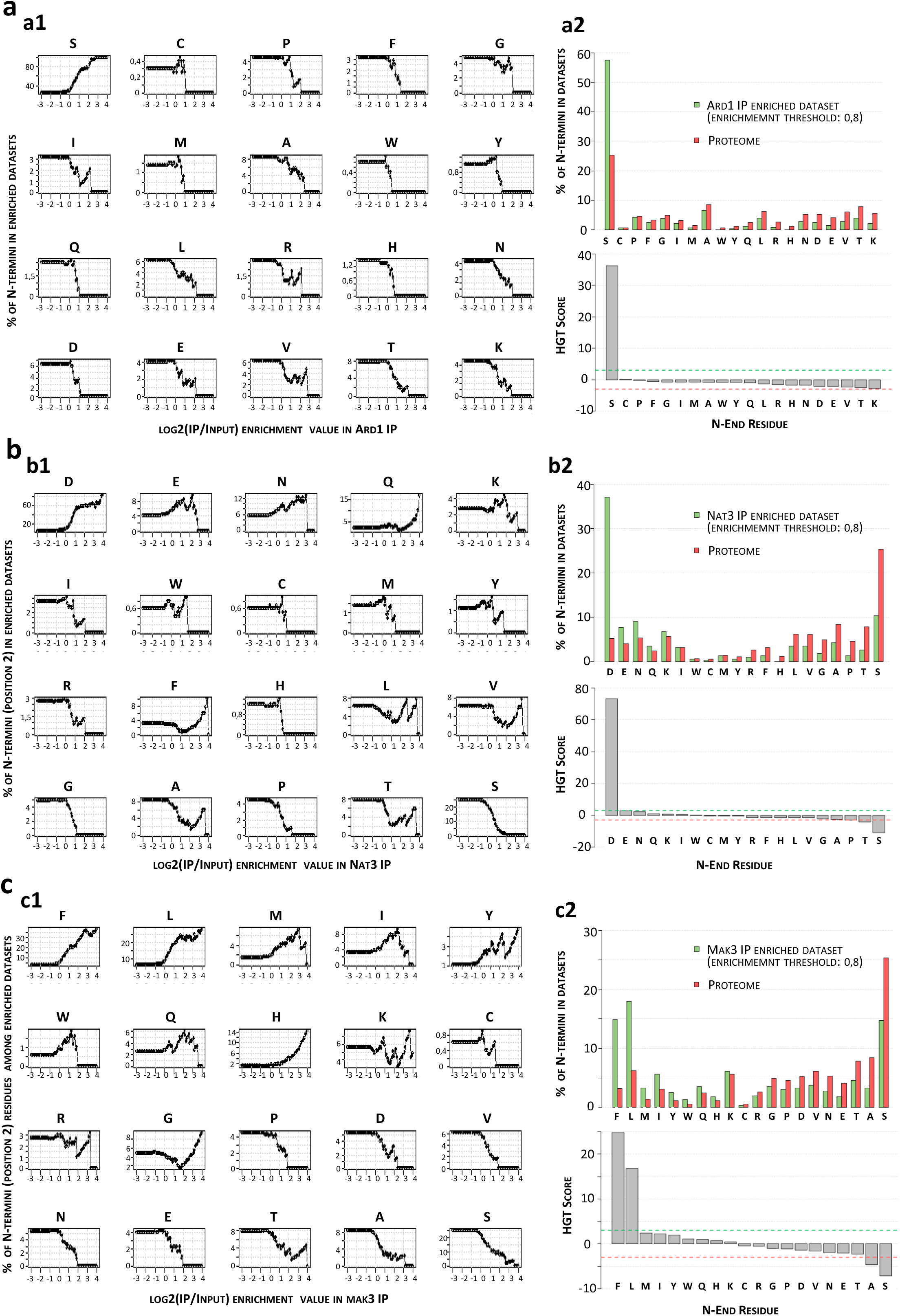
Analysis of the selectivity of NatA, NatB and NatC toward proteins N-termini using the sel-TRAP technology. Global analysis of mRNAs copurified after immunoprecipitations (IP) from cells expressing Ard1-PA (a), Nat3-PA (b) and Mak3-PA (c) allowed the indirect characterization of nascent protein chains translated by ribosomes associated respectively with NatA, NatB and NatC. For each cellular protein, the corresponding mRNA enrichment value, defined as IP/input log2(ratio), was calculated after microarray hybridizations of cDNAs reverse-transcribed from immunoprecipitation and input samples. Since NATs are known to target proteins according to the nature of their N-terminus, the left panels (a1, b1, c1) show the 20 curves representing, for all possible residues at position 2, the evolution of their proportion associated with the subsets of mRNAs detected with increasing enrichment values in IP experiments. After this analysis, an enrichment value of 0.8 was used as a threshold to define enriched mRNAs identified with sel-TRAP experiments of NatA, NatB and NatC. The percentages of each residue at position 2 among the identified targets were compared to those observed in the proteome (a2, b2, c2: upper panels) and the corresponding HGT scores were calculated (a2, b2, c2: lower panels) for statistical evaluation of significant differences. Dotted lines indicate the HGT score cutoff of 3 corresponding to a p-value obtained by a hypergeometric test equal to 10^−3^. This analysis of the sel-TRAP data revealed the specific co-translational selectivity of NatA for serine (S), NatB for aspartate (D), and NatC for phenylalanine (F) and leucine (L).

**Fig Sup 4:**
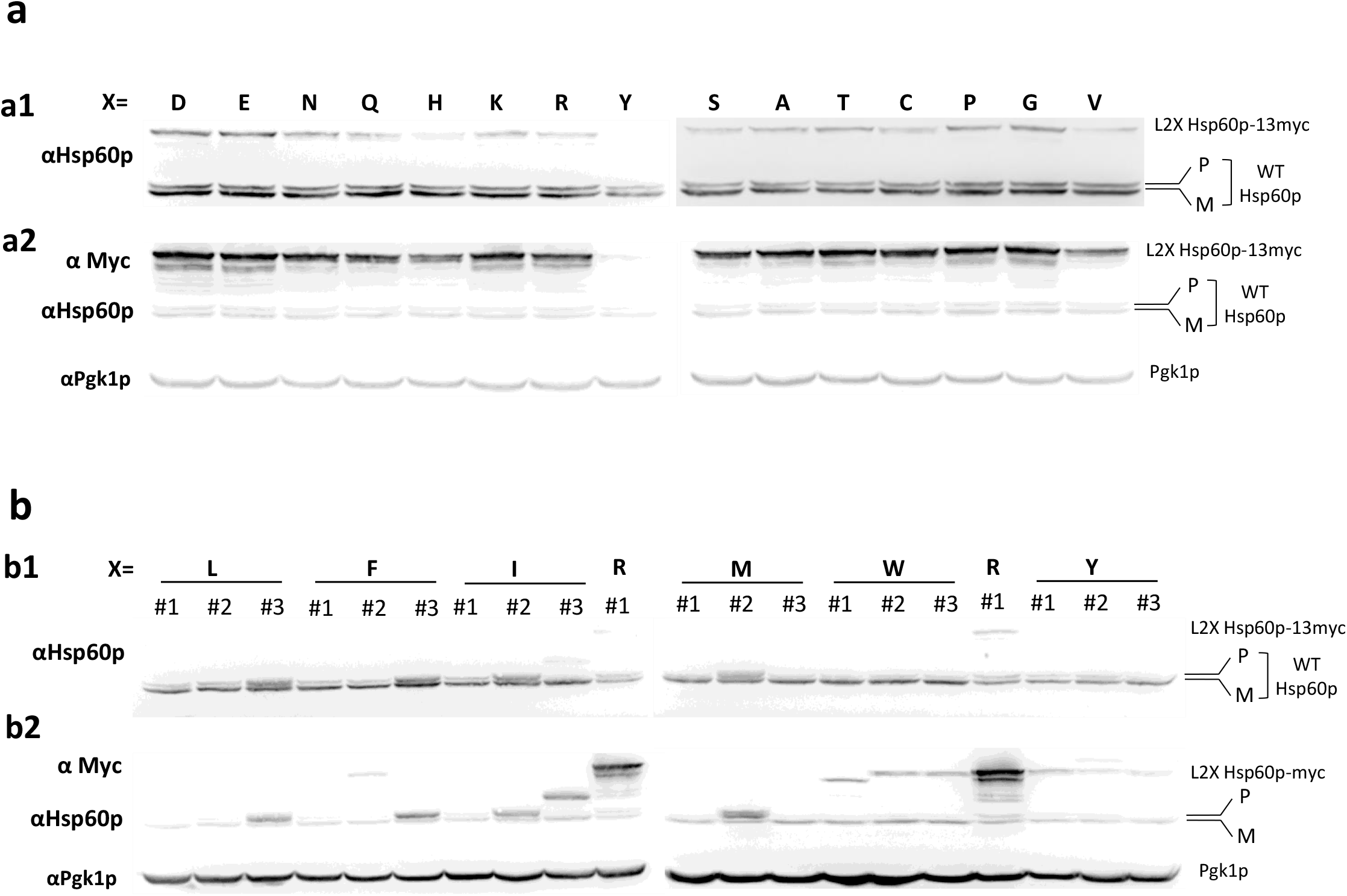
Western blot analysis of protein content in clones obtained after transformation with the diverse pHSPmyc(X) plasmids. Protein extracts were analyzed by SDS-PAGE followed by western blots using antibodies against myc epitope, Hsp60p, and Pgk1p. The position of the precursor and mature Hsp60p are indicated. (a) Analysis of clones transformed with plasmids that allow the rescue of the dominant negative toxicity of the Hsp60p−13myc protein (X=D, E, N, Q, H, K, R, Y, S, A, T, C, P, G, V, see Fig 5). (b) Analysis of clones transformed with plasmid expressing a toxic Hsp60p−13myc protein (X= L, F, I, M, W, see Fig 5) and of those obtained with the pHSP60(Y) for which a longer exposure time was necessary to detect the Hsp60p−13myc protein (see Fig. Sup3a).

## SUPPLEMENTARY METHODS

**Table S5:** Yeast strains.

| Strain | Genotype | Origin |
| --- | --- | --- |
| BY4741 | MATa; his3Δ1; leu2Δ0; met15Δ0; ura3Δ0 | Euroscarf (Brachmann et al,1998) |
| Ard1-PA | MATa; his3Δ1; leu2Δ0; met15Δ0; ura3Δ0, ARD1-ProtA-His5 | This Study*<sup>1</sup> |
| Nat3-PA | MATa; his3Δ1; leu2Δ0; met15Δ0; ura3Δ0, NAT3-ProtA-His5 |  |
| Mak3-PA | MATa; his3Δ1; leu2Δ0; met15Δ0; ura3Δ0, MAK3-ProtA-His5 |  |
| YPH499 | MATa ura3-52 lys2-801_amber ade2-101_ochre trp1-Δ63 his3-Δ200 leu2-Δ1 | Mehawej et al, PLoS Genet, 2014 |
| YPH499 pam16Δ-MAGN76D | MATa ura3-52 lys2-801_amber ade2-101_ochre trp1-Δ63 his3-Δ200 leu2-Δ2 pma16Δ + pmagMAT N76D |  |

\*<sup>1</sup>Strains tagged at their C terminus were obtained from BY4741 strain, after classical procedure of lithium acetate transformation, by homologous recombination with ProtA-His5 cassette amplified from pBXA (Rout et al., 2000 , provided by M. Rout, The Rockefeller, University, New York, NY)

**Table S6:**
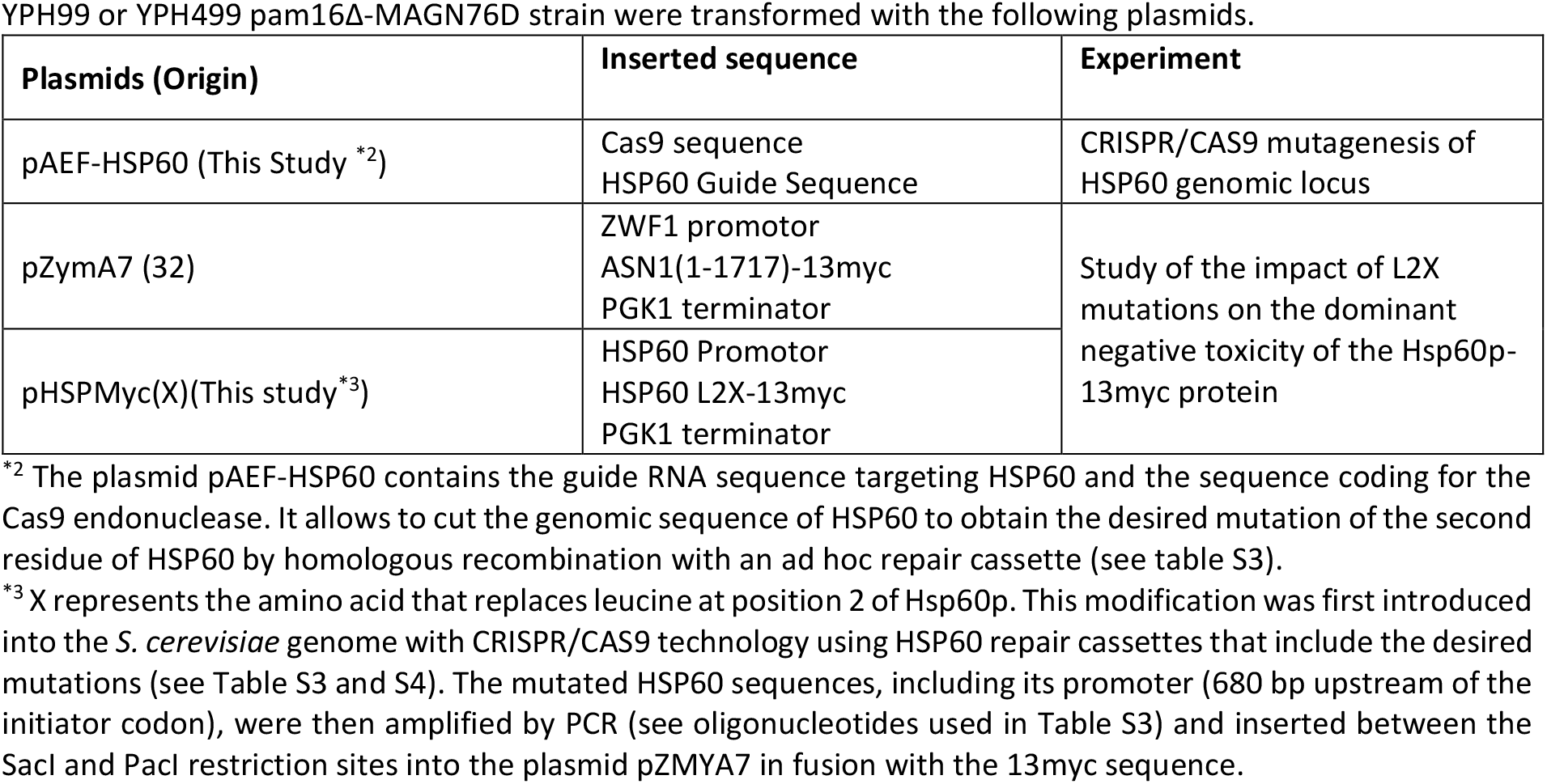
Plasmids and derived strains.

YPH99 or YPH499 pam16Δ-MAGN76D strain were transformed with the following plasmids.
| Plasmids (Origin) | Inserted sequence | Experiment |
| --- | --- | --- |
| pAEF-HSP60 (This Study *<sup>2</sup>) | Cas9 sequence<br>HSP60 Guide Sequence | CRISPR/CAS9 mutagenesis of HSP60 genomic locus |
| pZymA7 (32) | ZWF1 promotor<br>ASN1(1-1717)-13myc<br>PGK1 terminator | Study of the impact of L2X mutations on the dominant negative toxicity of the Hsp60p-13myc protein |
| pHSPMyc(X)(This study*<sup>3</sup>) | HSP60 Promotor<br>HSP60 L2X-13myc<br>PGK1 terminator |  |

\*<sup>2</sup> The plasmid pAEF-HSP60 contains the guide RNA sequence targeting HSP60 and the sequence coding for the Cas9 endonuclease. It allows to cut the genomic sequence of HSP60 to obtain the desired mutation of the second residue of HSP60 by homologous recombination with an ad hoc repair cassette (see table S3).
\*<sup>3</sup> X represents the amino acid that replaces leucine at position 2 of Hsp60p. This modification was first introduced into the *S. cerevisiae* genome with CRISPR/CAS9 technology using HSP60 repair cassettes that include the desired mutations (see Table S3 and S4). The mutated HSP60 sequences, including its promoter (680 bp upstream of the initiator codon), were then amplified by PCR (see oligonucleotides used in Table S3) and inserted between the SacI and PacI restriction sites into the plasmid pZMYA7 in fusion with the 13myc sequence.

**Table S7:**
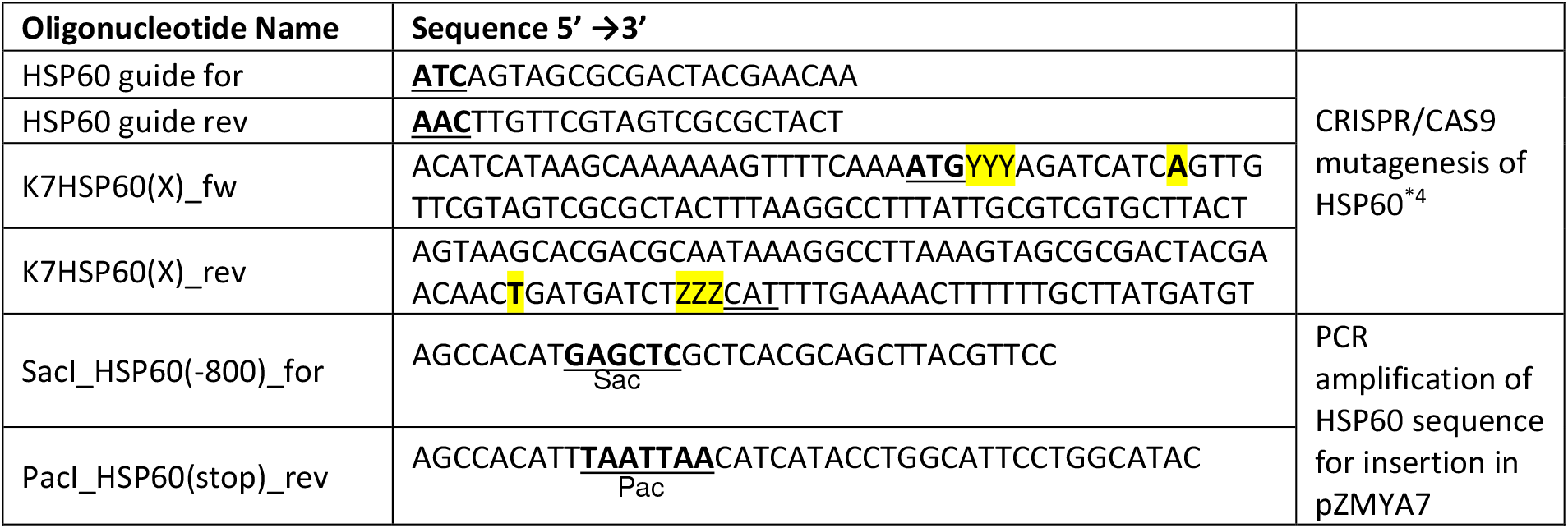
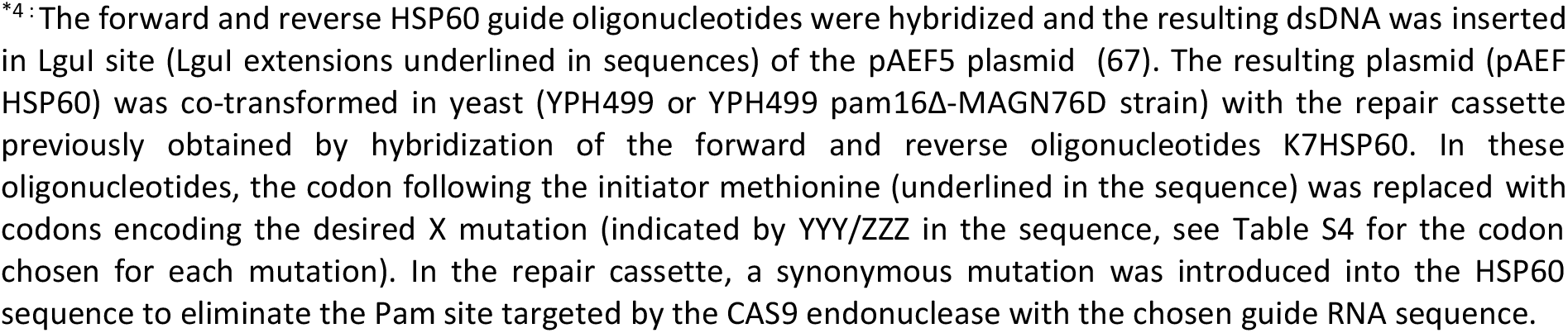
Oligonucleotides.

**Table S8:**
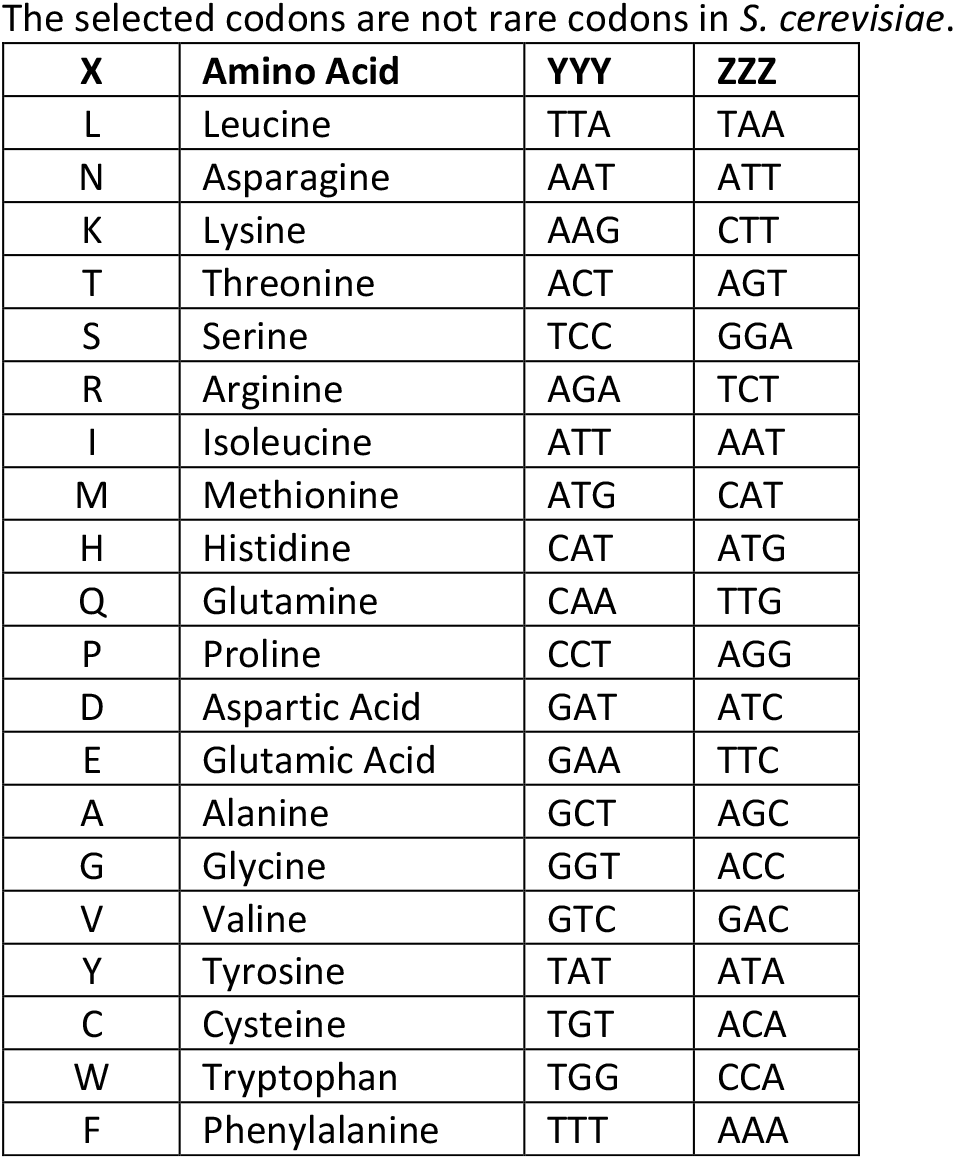
Codon used in the reparation cassette to encode the desired X mutation.

The selected codons are not rare codons in *S. cerevisiae*.
| X | Amino Acid | YYY | ZZZ |
| --- | --- | --- | --- |
| L | Leucine | TTA | TAA |
| N | Asparagine | AAT | ATT |
| K | Lysine | AAG | CTT |
| T | Threonine | ACT | AGT |
| S | Serine | TCC | GGA |
| R | Arginine | AGA | TCT |
| I | Isoleucine | ATT | AAT |
| M | Methionine | ATG | CAT |
| H | Histidine | CAT | ATG |
| Q | Glutamine | CAA | TTG |
| P | Proline | CCT | AGG |
| D | Aspartic Acid | GAT | ATC |
| E | Glutamic Acid | GAA | TTC |
| A | Alanine | GCT | AGC |
| G | Glycine | GGT | ACC |
| V | Valine | GTC | GAC |
| Y | Tyrosine | TAT | ATA |
| C | Cysteine | TGT | ACA |
| W | Tryptophan | TGG | CCA |
| F | Phenylalanine | TTT | AAA |

**Table S9:**
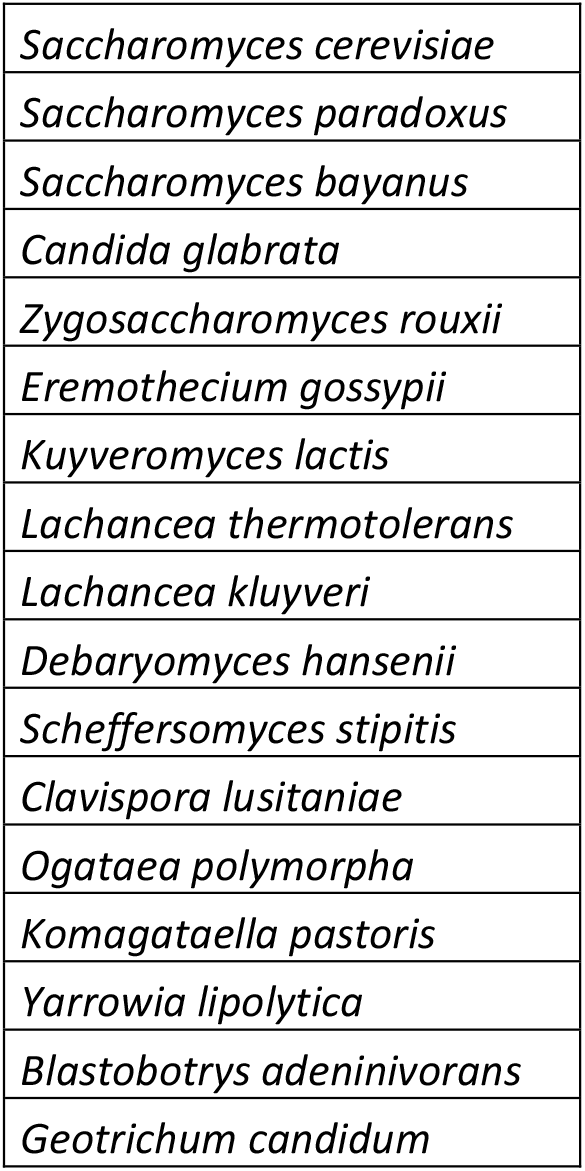
List of the yeast species used for the genomic comparative studies.

## GO reduction algorithm

### Definitions

Let *G_i_* be the ensemble of genes from the protein-coding fraction 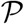 associated with the GO term *GO_i_*, such that *G_i_* = 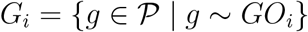, where *g* is a gene and the symbol ∼ denotes the association between a gene and a GO term. We define 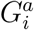 as the subset of genes from *G_i_* whose resulting proteins display the amino acid *a* in position 2. We evaluate the representativity of a GO term with respect to *a* according to the following score,

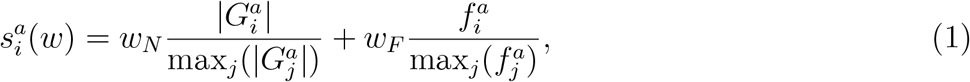

where 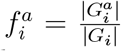 is the frequency of occurrence of *a* in the second position of the proteins resulting from the genes included in *G_i_*, and *w_N_* and *w_F_* are weights controlling the relative contributions of the two terms such that *w_N_* + *w_F_* = 1. The first term reflects how well the GO term *GO_i_* covers the proteins displaying *a* in position 2. The GO term associated with the maximum number of such proteins has the maximum value of 1. The second term reflects how specific the GO annotation is for the amino acid *a*. The lower the variability of amino acids in position 2 among the proteins associated with a given GO term, the higher this value.

### Initialization: Pre-selection of GO terms

We start from 20 sets of pre-selected GO terms corresponding to the 20 amino acids. For each amino acid *a*, the *N_a_* selected GO terms are those displaying an enrichment for *a* with a p-value *p* − *val* ≤ 10^−4^, and also for which 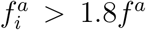, where *f^a^* is the frequency of occurrence of *a* at position 2 in the proteome. Notice that some sets may be empty.

### First step: identifying the most inclusive and selective GO terms for each amino acid

The goal of this first step is to define, for each amino acid, a minimal set of GO terms representative of the full pre-selected set, both in terms of coverage and of specificity. We first order the GO terms according to the score expressed in Eq. 1, from the highest to the lowest score. We set the weight *w_N_* = *w_F_* = 0.5 so that coverage and specificity contribute equally to the score. Then, we apply the following Algorithm 1, independently for each amino acid *a*. The algorithm considers the *N_a_* GO terms pre-selected for *a*, from the highest-scored one to the lowest-scored one, and decides whether each term should be retained or discarded based on its gene overlap with the previously retained terms. At each iteration of the algorithm, we require that the GO term considered shares less than α = 40% of its associated genes with the current GO term list. If this criterion is not met, then the GO term is disregarded.

### Second step: reducing redundancy at the global level while prioritizing coverage

While the first step treats each amino acid independently from the others, this second step aims at reducing redundancies across all amino acids. More precisely, for each amino acid *a*, we look for the GO terms not included in its *bestNGO* list but present in the *bestNGO* list of at least one other amino acid *b* and displaying an enrichment for *a* with a p-value *p* − *val* ≤ 10^−3^. We extend the *bestNGO* list of *a* with these GO terms. They are of interest because they display high enrichments for several amino acids and they potentially cover more genes than the terms selected in the previous step. To allow for them to be retained, and potentially replace GO terms with lower coverage, we again apply Algorithm 1 for |each amino acid setting *w_N_* = 1 and *w_N_* = 0.

#### Algorithm 1 Redundancy reduction applied to a list of pre-selected GO terms

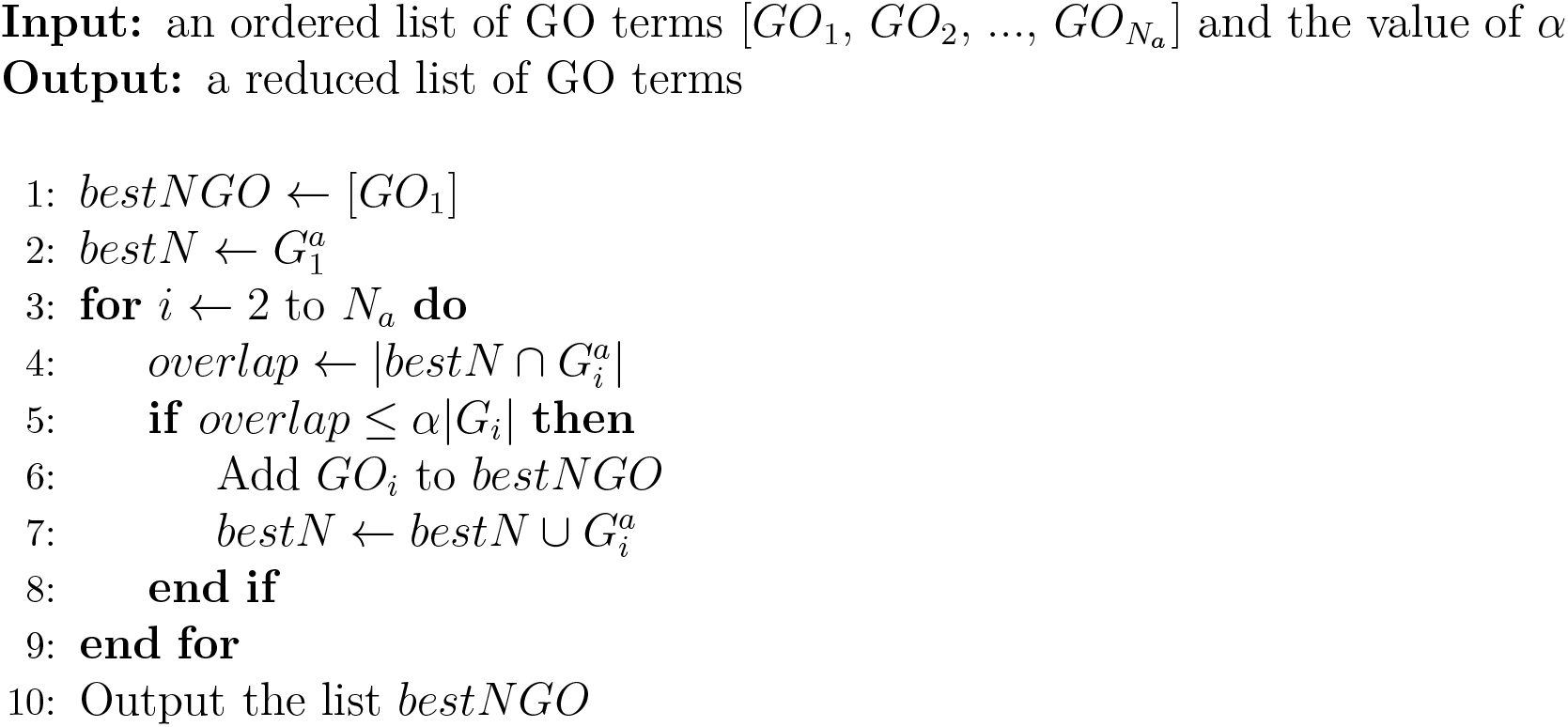

#### Algorithm 2 Rescue of GO terms from a pre-filtered list

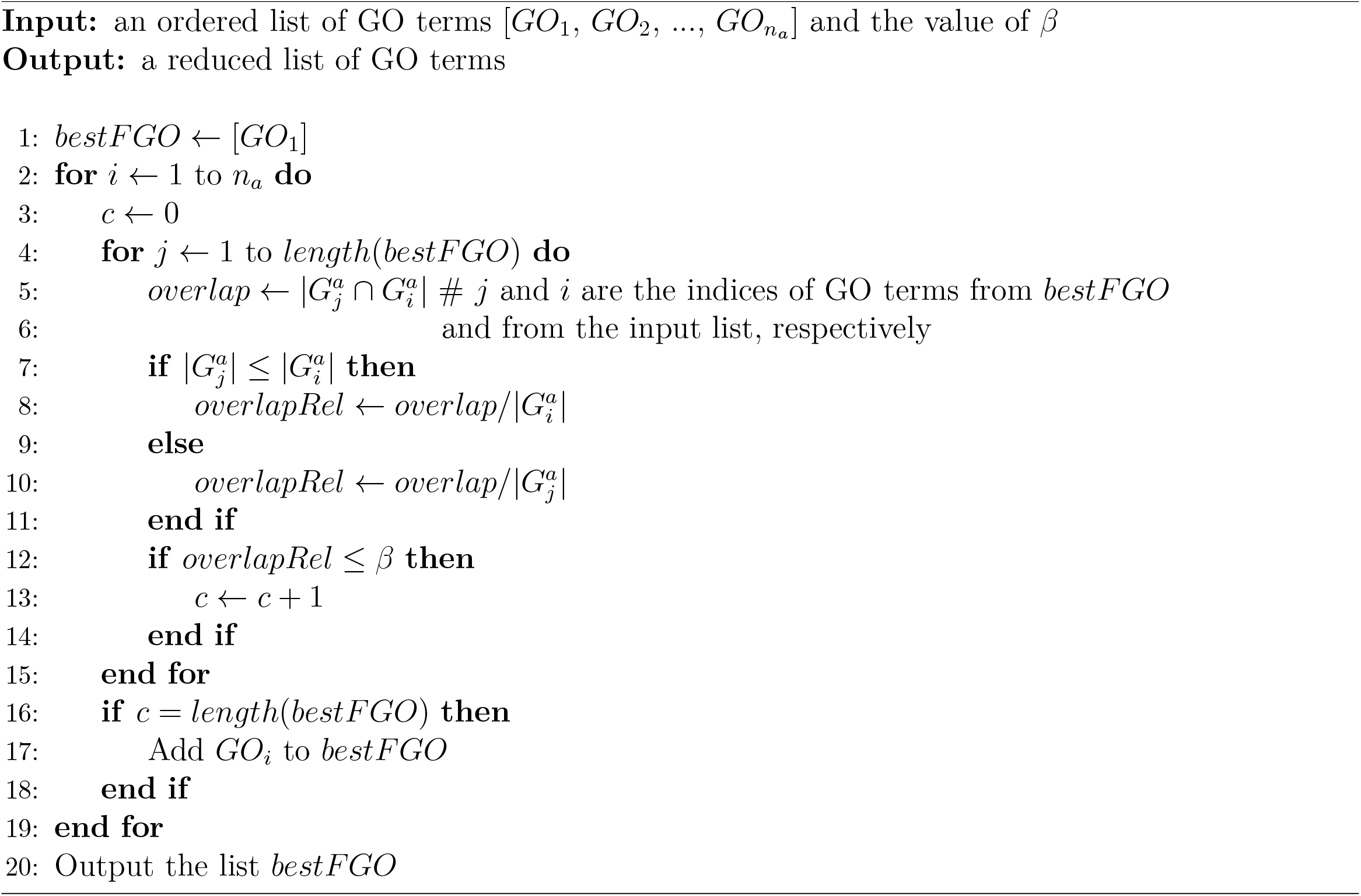

### Third step: rescuing highly selective GO terms

This goal of this third step is to rescue the GO terms displaying a very high selectivity for a particular amino acid. Here, for each amino acid *a*, we start from the *n_a_* GO terms pre-selected in the initialization phase but not retained in the *bestNGO* list. We order them according to the score expressed in Eq. 1, from the highest to the lowest score, with the weights set to *w_N_* = 0 and *w_F_* = 1. The rationale is to prioritize the most selective GO terms. Then, we apply the following Algorithm 2. At each iteration, we require for a GO term to be retained that it shares less than β = 30% of its associated genes with each one of the GO terms in the current list.

### Fourth step: reducing redundancy at the global level while prioritizing specificity

Similarly to what is done in the second step, the goal of this fourth step is to reduce redundancies across all amino acids. We consider the *nF* GO terms appearing in at least one of the *bestFGO* lists defined in the previous step. For each of these GO terms, we identify the amino acids for which it displays an enrichment with a *p* − *val* ≤ 10^−4^. This operation leads to *nF* potentially overlapping subsets of amino acids. The subsets with only one member are discarded. We encode this information in a graph where the nodes are the 20 amino acids and the set of edges is defined from the subsets.

#### Algorithm 3 Build a graph representing the overlaps between GO terms enriched for a set of amino acids

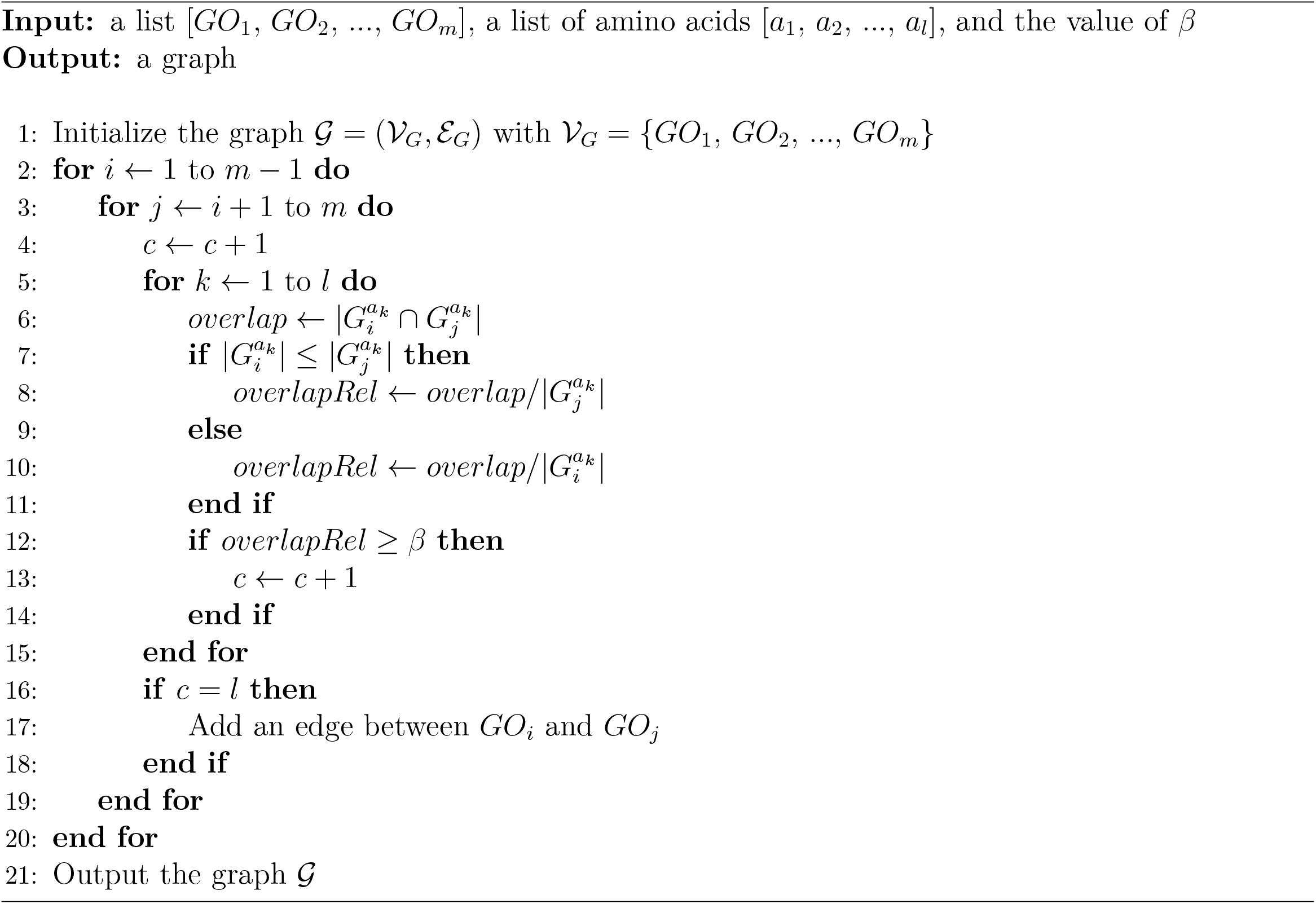

More specifically, any pair of nodes is linked by an edge if the corresponding amino acids were found in the same subset at least once. We further define groups of amino acids as the connected components of the graph. Each group {*a*_1_, *a*_2_, …, *a_l_*} is associated with a set of GO terms {*GO*_1_, *GO*_2_, …, *GO_m_*} coming from the *bestFGO* lists. We then build a graph encoding the gene overlaps between these GO terms by applying Algorithm 3. For each connected component in the graph, we determine a representative GO term *GO_imax_* as the most selective one for the amino acid group. Formally, we determine arg 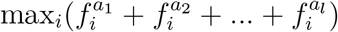. The final operation consists in replacing all the GO terms appearing in the *bestFGO* lists defined for the amino acids *a*_1_, *a*_2_, …, *a_l_* by their representative *GO_imax_*.

### Mass spectrometry analysis

Digestion was performed overnight at 37°C in the presence of 12.5 μg/ml of sequencing grade trypsin (Promega, Madison, Wi, USA). Peptides mixtures were analyzed by a Q-Exactive Plus coupled to a Nano-LC Proxeon 1000 (both from Thermo Scientific). Peptides were separated by chromatography using the following conditions: Acclaim PepMap100 C18 pre-column (2 cm, 75 μm i.d., 3 μm, 100 Å), Pepmap-RSLC Proxeon C18 column (50 cm, 75 μm i.d., 2 μm, 100 Å), 300 nl/min flow rate, a 98 min gradient from 95 % solvent A (water, 0.1 % formic acid) to 35 % solvent B (100 % acetonitrile, 0.1% formic acid) followed by column regeneration, giving a total time of 120 minutes. Precursor peptides were analyzed in the Orbitrap cell in positive mode, at a resolution of 70,000, with a mass range of *m/z* 375−1500 and an AGC target of 3.10^6^. MS/MS data were acquired in the Orbitrap cell in a Top20 data-dependent mode with a dynamic exclusion of 30 seconds. Fragments were obtained by Higher-energy C-trap Dissociation (HCD) activation with a collisional energy of 27% and a quadrupole isolation window of 1.4 Da. The Orbitrap cell was set at a resolution of 17,500, *m/z* 200−2000 and an AGC target of 2.10^5^. Peptides with unassigned charge states or monocharged were excluded from the MS/MS acquisition. The maximum ion accumulation times were set to 50 ms for MS acquisition and 45 ms for MS/MS acquisition.

Data were processed with Proteome Discoverer 2.2 software (Thermo Fisher scientific, San Jose, CA) coupled to an in-house Mascot search server (Matrix Science, Boston, MA; version 2.5.1). MS/MS spectra were searched against the SwissProt protein database release 2017_09 with the *Saccharomyces Cerevisiae (baker’s yeast)* taxonomy and a maximum of 2 missed cleavages. Precursor and fragment mass tolerances were set to 6 ppm and 0.02 Da respectively. The following post-translational modifications were included as variable: Acetyl (Protein N-term), Oxidation (M), Phosphorylation (STY). Spectra were filtered using a 1% FDR with the percolator node.

## Notes

### Competing Interest Statement

The authors have declared no competing interest.

### Summary of Updates

Correction of the order of the paragraphs in the results section

